# Chronology of motor-mediated microtubule streaming

**DOI:** 10.1101/505669

**Authors:** Arvind Ravichandran, Özer Duman, Masoud Hoore, Guglielmo Saggiarato, Gerard A. Vliegenthart, Thorsten Auth, Gerhard Gompper

## Abstract

We introduce a computer-based simulation model for coarse-grained, effective motor-mediated interaction between microtubule pairs to study the time-scales that compose cytoplasmic streaming. We characterise microtubule dynamics in two-dimensional systems by chronologically arranging five distinct processes of varying duration that make up streaming, from microtubule pairs to collective dynamics. The structures found were polarity sorted due to the propulsion of antialigned microtubules. This also gave rise to the formation of large polar-aligned domains, and streaming at the domain boundaries. Correlation functions, mean squared displacements, and velocity distributions reveal a cascade of processes ultimately leading to microtubule streaming and advection, spanning multiple microtubule lengths. The characteristic times for the processes span over three orders of magnitude from fast single-microtubule processes to slow collective processes. Our approach can be used to directly test the importance of molecular components, such as motors and crosslinking proteins between microtubules, on the collective dynamics at cellular scale.

## I. INTRODUCTION

The vigorous motion of the intracellular fluid, known as cytoplasmic streaming, is caused by cytoskeletal filaments and molecular motors. In *Drosophila* oocytes this cellular-scale fluid motion, which occurs over multiple time scales, is responsible for efficient mixing of ooplasm and nurse-cell cytoplasm, for long-distance transport of intracellular material and for proper patterning of the oocyte [1–3]. Although cytoplasmic streaming is known for centuries [4] and kinesin-1 molecular motors and microtubules (MTs) have been identified as the components responsible for ooplasmic streaming [1, 2, 5, 6], there is considerable debate about the aetiological mechanisms for force generation. Namely, the constituent events, their order of occurrences, and their characteristic durations, which ultimately give streaming are not understood. Some studies suggest that streaming is caused by the hydrodynamic entrainment of motor-transported cargos [7, 8], others that it is due to the motor-mediated sliding of adjacent MTs [3, 9, 10].

Motor-mediated MT sliding occurs because molecular motors crosslink adjacent MTs and use ATP (adenosine triphosphate) molecules as fuel to “walk” on them unidirectionally in the direction of MT polarity [11]. This leads to significantly different active dynamics of MT pairs that are polar-aligned and antialigned [12–14]: Motors that crosslink polar-aligned MTs hold the polar-aligned MTs together, generating an effective attraction [14]. Active motors crosslink and slide antialigned MTs. The motors thus act as force dipoles that break nematic symmetry in MT solutions. In the absence of permanent crosslinkers, which are known to render the active network contractile [15], this ultimately can cause largescale flows in the cytoskeleton [3, 9]. Several approaches to analyse the collective motion, such as the displacement correlation function or the analysis of velocity distributions, are inspired by studies on collective motion of self-propelled agents [16–20].

*In vivo*, individual MTs are stationary most of the time before suddenly undergoing a burst of long-distance travel with velocities reaching *≈* 10 *µ*m/s [9]. Also, fluorescence microscopy has shown the formation of long extended arms for an initially circular photoconverted area [9]. A possible mechanism for such a behaviour is illustrated in Fig. 1: In the absence of active motor stresses, MTs in polar-aligned bundles diffuse slowly. When they encounter antialigned MTs, they can be actively and rapidly transported away from a polar-aligned bundle to another polar-aligned bundle where they again exhibit slow diffusive behaviour. These transitions between slow-bundling motion and fast-streaming bursts can give rise to Levy flight-like MT dynamics [21].

**Figure 1.**
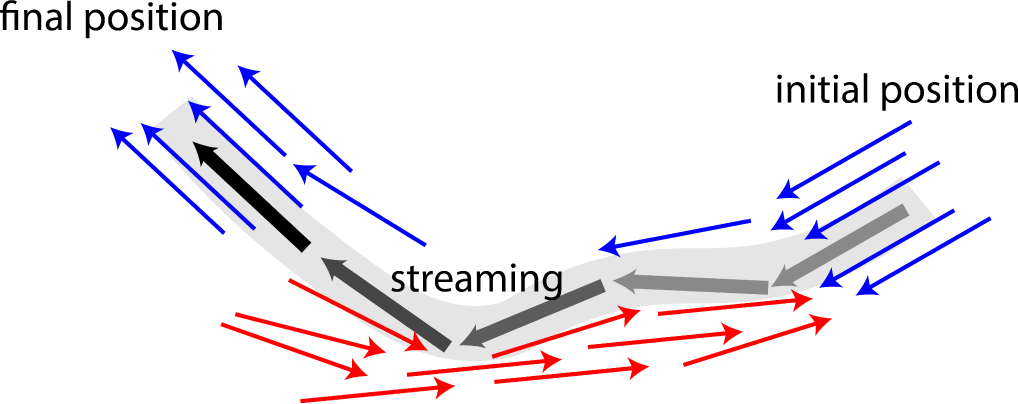
Schematic illustrating MT bundling and streaming. Polar-aligned MTs are coloured blue, and antialigned MTs are coloured red. The grey/black MT is transported from its initial position (grey), in one polar-aligned bundle, to its final position (black), to another polar-aligned bundle, via a stream.

*In vitro* and *in silico*, MT-motor model systems allow a systematical study of the interactions between the basic components of the cytoskeleton. The systems often contain only a small number of different components and are usually also restricted in other respects, such as a reduced dimensionality or a lack of polymerization and depolymerization of the filaments. For example, a very well studied model system contains MTs and kinesin complexes at an oil-water interface [22, 23], where poly(ethylene glycol)-induced depletion interaction both keeps the MTs at the interface and induces bundle formation. In contrast, coarse-grained computer simulations in 2D are used to study both structure formation on the bundle scale [23] as well as single-filament dynamics, e.g. to quantify the effect of motor properties or of the presence of passive crosslinkers [12–15]. Because such model systems are simpler than the cytoskeleton in biological cells, they are especially suited to study specific mechanisms in detail.

Cytoplasmic streaming is a complex multi-scale phenomenon that cannot be fully understood using antialigned filaments alone. The importance of a specific mechanism can only be studied *in vivo* for a specific system. This so-called ‘top-down’ approach has been remarkably successful in describing streaming in the aquatic alga *Chara coranilla* [24] and *Dropsophila* oocytes [25]. In the former case, theoretical models have showed the importance of coupling hydrodynamic entrainment and microfilament dynamics to capture pattern formation relevant for streaming. In the latter case, simulations mimicking streaming in *Drosophila* oocytes have emphasised the importance of cortical MT nucleation in anteroposterior axis definition. It was shown that nucleation of MTs from the periphery is important to induce cytoplasmic flow patterns and to localise mRNAs in specific areas of the cell.

Microtubule-motor systems are intrinsically out of equilibrium, which has been shown for example by monitoring the dynamics of motors that walk along MTs [26] and by the violation of the fluctuation-dissipation theorem for tracer particles embedded into acto-myosin systems [27]. Therefore, simulation approaches for systems of passive MTs at equilibrium have to be augmented with motor activity. Brownian dynamics simulations with MTs and explicit motors have been used to study network contractility [15], polarity-sorting and stress generation at high MT densities [12], and persistent motion of active vortices in confinement [28]. Recent simulations for the defect dynamics in extensile MT systems have been performed on the coarse-grained level of MT bundles [23].

We employ a ‘bottom-up’ approach, where we study MT streaming induced by MT sliding using a model system. In order to characterize the dynamics in the system, we use a coarse-grained model to investigate whether a purely polarity-dependent MT-MT sliding mechanism, in the absence of any hydrodynamic forces, can be sufficient to capture large-scale streaming in bulk. We identify five distinct processes that comprise streaming with their characteristic times for various MT activities and surface fractions: 1) motor-driven MT sliding, 2) polarity-inversion, 3) maximal activity, 4) collective migration, and 5) rotation. Although various experimental studies have provided high spatial resolution to describe streaming phenomena [1], MT dynamics for streaming is still poorly understood. Inspired by the biological mechanism for MT-MT sliding, we use computer simulations to provide a novel temporal perspective into streaming for a wide range of time scales, which has not been achieved so far due to limitations of experimental techniques.

In order to capture cellular-scale dynamics in computer simulations, modelling individual motors along with MTs, although done before in several studies [12, 13, 27, 29–32], can prove to be unwieldy due to the wide ranges of length and time scales involved. The sizes of individual kinesin molecules that crosslink and slide MTs are three orders of magnitude smaller than that of the cells within which they bring about large-scale dynamics. Also, there is a large disparity between the residence time of a cross-linking motor (10 seconds) [33], and the characteristic time scale of motor-induced MT streaming or pattern formation in active gels (1 hour) [3, 9, 23, 34]. In order to capture motor-induced cellular-level phenomena, such as organelle distribution, cytoplasmic streaming, and active cytoskeleton-induced lipid bilayer fluctuations, a coarse-grained description of cytoskeletal activity seems therefore appropriate.

In this work, we use a model of MTs interacting via the polarity dependent interaction force to study the non-equilibrium structure and dynamics of MTs, and the phenomena of MT streaming, as a function of the MT density and the probability that a motor crosslinks two MTs. The model is studied in two dimensions by Langevin dynamics simulations. The coarse-grained MT-motor model is explained in Sec II, the simulation results are presented in Sec III, and we discuss the results and summarise our conclusions in Sec IV.

## II. MICROTUBULE-MOTOR MODEL

### A. Active-Microtubule Suspensions

Coarse-grained and continuum approaches are successfully applied to study cytoskeletal-motor systems. A well-developed model and simulation package is Cytosim that can be used to simulate flexible filaments together with further building blocks that, for example, act as nucleation sites, bind filaments together, and induce motility or severing [35]. It has been applied to study–among other processes in the cell–meiosis [36], mitosis [37], and centrosome centering [38]. A different model that includes MT flexibility, MT polymerization and depolymerization, explicit motors, and hydrodynamics has recently been applied to study mitosis [39, 40]. Because the MTs are mostly radially oriented, steric interactions between MTs can be neglected and have not been taken into account. However, MT-MT repulsion is important to obtain nematic order at high MT densities, an essential ingredient for the bottom-up model systems containing suspensions of MTs and kinesins [22]. Our model includes MT flexibility, effective-motor potentials, and excluded-volume interactions. Polymerization and depolymerization does not occur in the model system and is not taken into account. Effective-motor models in general aim to reduce the computational effort to efficiently study large systems [41, 42]. Including hydrodynamic interactions using a particle-based approach is straightforward [43–45], but beyond the scope of this paper. The model for the MTs is introduced in section II B, the active orientation-dependent motor potential *U*_mot_ is introduced in section and II C. The Langevin Dynamics used to simulate our system is described in section II D, and typical parameter values are discussed in section II E.

### B. Microtubules

In our two-dimensional model, MTs are modelled as impenetrable, semi-flexible filaments of length *L*, thickness *σ*, and aspect ratio *L/σ*, under periodic boundary conditions. Each of the *N* filaments in the system is discretised into a chain of *n* beads with diameter *σ* that are connected by harmonic bonds. The configurational potential,

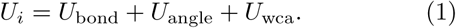

is the sum of passive potentials, *i.e.* the spring potential *U*_bond_ between adjacent beads, the angle potential *U*_angle_ between adjacent bonds, and the volume exclusion *U*_WCA_ between MTs.

The bond energy,

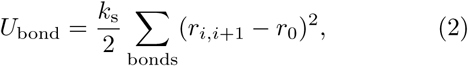

acts between adjacent beads of the same MT. Here, *k*_s_ is the bond stiffness, *r*_0_ = *σ/*2 is the equilibrium bond length, and *r*_*i,i*+1_ = |**r**_*i,i*+1_ | is the distance between adjacent beads *i* and *i* + 1, which make up the MT. *U*_angle_ is the bending energy, which is calculated using the position of three adjacent beads,

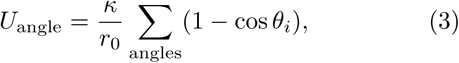

that make up the angle *θ*_*i*_ [46]. It acts between all groups of three adjacent beads that make up the same MT. The bending modulus *κ* of the filament determines its persistence length *𝓁*_p_ = *κ/k*_B_*T*.

MT bead pairs that are not connected by harmonic springs interact with each other via the repulsive Weeks-Chandler-Andersen (WCA) potential [47],

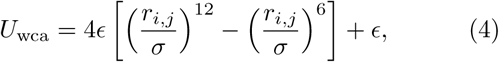

with interaction cutoff *r*_cut_ = 2^1^*/*^6^*σ*.

### C. Effective motor potential

Various theoretical studies have strived to circumvent short time and length scales involved in cytoskeletal dynamics, such as diffusion and active motion of individual motors [41, 42, 48–53]. For example, in the phenomenological flux-force model the motion of MTs in one dimension occurs solely due to the orientation of neighbouring MTs [54, 55]. Many two-dimensional models, where MTs are modelled as stiff, polar rods of equal length, take motors into account using a Maxwellian model of inelastic interactions between the rods [41, 42, 50, 51]. These probabilistic collision rules result in the alignment of rods. Although these models capture the selforganization of MT-motor mixtures into stable patterns of vortices, asters, and smectic bundles, the collision rule does not reproduce the sliding of antialigned MTs described in Fig. 2.

**Figure 2.**
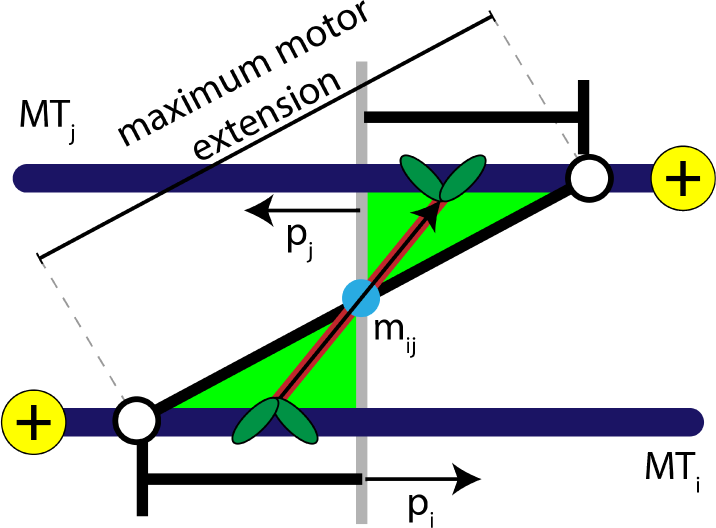
Schematic explaining the conditions that satisfy the antialigned motor potential. The vectors, **p**_*i*_, **p**_*j*_, and **m**_*ij*_, represent the unit orientation vectors of MT *i*, MT *j*, and the motor vector that crosslinks the beads of adjacent MTs, respectively. The white circles represent the maximum extension of motors between the two MTs.

Sliding of antialigned MTs due to kinesin motors has been identified as key ingredient for cytoplasmic advection *in vivo* [3, 9, 10]. Instead of modelling individual motors, in our model MT motion manifests itself as a result of a distribution of motors in an ensemble of orientations between neighbouring MT pairs. Hence, we coarse-grain MT-motor interactions using an effective motor potential that gives a contribution to *U*_mot_. A motor bond can form when the crosslinked beads are antialigned, *i.e.*, the angle that a motor bond vector **m**_*ij*_ makes with the unit orientation vector is acute on both MTs simultaneously, see Fig. 2,

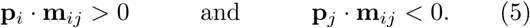

Here the orientation vector assigned to bead *i* on an MT is **p**_*i*_ = (**r**_*i*+1_ *-***r**_*i*_)*/* |**r**_*i*+1_ *-***r**_*i*_ |, and the extension of a motor that crosslinks MTs *i* and *j* is **m**_*ij*_(*s*_*i*_*, s*_*j*_) = **r**_i_(*s*_*i*_) *-***r**_j_(*s*_*j*_), with the motor heads bound at the positions *s*_*i*_ and *s*_*j*_ along the contour of the MTs. This is similar to the activity-inducing scenario a dimeric or tetrameric motor [14] encounters when it crosslinks a pair of antialigned MTs, *i.e.*, the motor arms are oriented towards the + direction of either crosslinked MT.

Each effective motor is a harmonic spring of equilibrium bond length *d*_eq_ = *σ* and stiffness *k*_m_ that exists for one simulation time step [56]. The system is inherently out of equilibrium because the motor bonds occur dependent on the relative orientation of neighboring MTs, and exist and exert forces only for short times, mimicking the ratchet model for molecular motors [57]. The potential for a motor with extension *m*_*ij*_ = |**m**_*ij*_| is

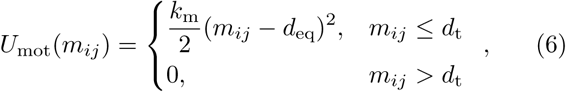

and the motor binding rate is

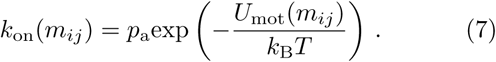

Here, *p*_a_ controls the probability that an antialigned motor binds [58]. Motors bind only for extensions *m*_*ij*_ *< d*_*t*_ = 2*σ*, when *k*_on_*/p*_a_ *>* exp(*-*1*/*2). This also corresponds well to the experimentally measured length of a kinesin motor [59]. This motor model described here for two dimensions is analogous to the phenomenological model for one dimension described in Ref. [48]; these onedimensional calculations show that the relative velocity between two antialigned MTs is a linear function of *p*_a_ and *k*_m_. Similarly, a kinesin-5 induced effective torque between MTs has been calculated to study forces in the mitotic spindle [60].

### D. Langevin Dynamics

The motion of the beads is described by the Langevin equation,

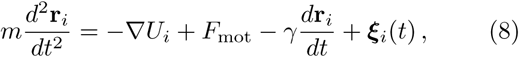

where **r**_*i*_ is the position of bead *i*, *m* is the mass of a bead, *γ* is the friction coefficient of the solvent for bead motion, *F*_mot_ = *−∇U*_mot_ is the active motor force and ***ξ***_*i*_ is the Gaussian-distributed thermal force. The friction coefficient can be estimated using the Stokes friction *γ* = 6*πηR* for a spherical particle with radius *R* in a solvent with viscosity *η*. The thermal forces ***ξ***_*i*_ have ⟨ ***ξ***_*i*_ ⟩ = 0 and, from the fluctuation-dissipation theorem,

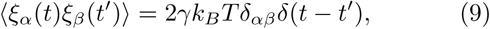

where *k*_*B*_ is the Boltzmann constant, T is the temperature, and *ξ*_*α*_(*t*) is the *α*-th component of the vector ***ξ***_*i*_(*t*).

Langevin dynamics simulations allow the use of larger time steps compared with Brownian dynamics without a particle mass. The friction constant *γ* and bead mass *m* are chosen such that the center-of-mass motion of passive MTs at the same density is diffusive at length scales larger than a fraction of the MT length and at time scales *τ/τ*_*R*_ *≥* 0.01 (see SI), such that passive MTs only move ballistically at times shorter than the relevant times.

Video 1. Simulation video of MT-effective motor system. MTs are coloured based on their orientation according to the colour legend given in Fig. 3(a). The video is recorded over a duration of 100*τR*.

**Figure 3.**
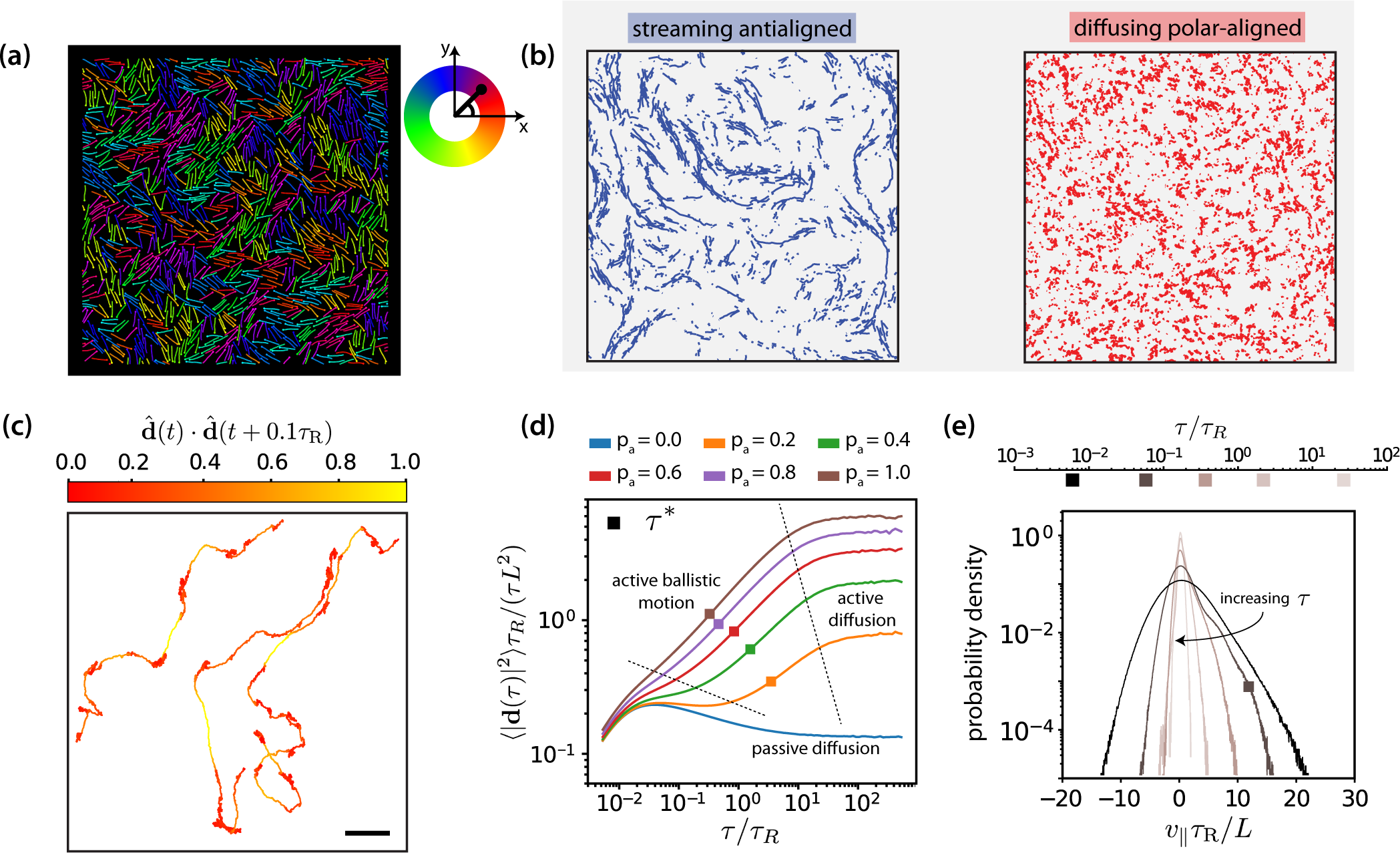
(a) Simulation snapshot of MTs organised by effective motors. MTs are coloured based on their orientation according to the colour legend on the right. See corresponding Video 1. (b) Trajectories of MTs within a time window of 1.2 *τR* separated based on the antialigned and polar-aligned categories. See corresponding Video 2. (c) Plots of the trajectory of three selected MTs coloured based on the correlation of adjacent steps in their velocity. The entire trajectory is for a time window of 300*τR*. 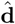 is the unit vector of MT displacement. The fast-streaming and slow-diffusion modes correspond with the yellow and red parts of the trajectories respectively. See corresponding Video 3. (d) MSD/lag time for various levels of activity *p*_a_ and MT density *ϕ* = 0.3. The time scale of maximal activity, *τ*^*\**^, calculated from the time of maximal *v*_‖_ skew is indicated by the squares on the curves. (e) Histogram of parallel velocity for various *τ*. The curve closest corresponding to the time scale of maximal activity, *τ*^*\**^, is indicated with a box marker. All figures are for *ϕ* = 0.3. (a), (b), (c) and (e) are for *p*_a_ = 1.0.

Video 2. Trajectories of MTs within a time window of 1.2*τR* separated based on the antialigned (left) and polar-aligned (right) categories. The video is recorded over a duration of 100*τR*.

The simulation package LAMMPS has been employed to perform the simulations [61].

### E. System parameters

Each simulation consists of *n*_f_ = 1250 semiflexible filaments with aspect ratio 10, each made up of *n*_b_ = 21 overlapping beads, which reduces the friction of the otherwise corrugated MTs [64, 70]. The MT surface fraction 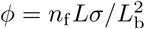 is controlled by adjusting the box size *L*_b_. Our effective-motor model is a coarse-grained model and individual effective motors in simulation may not represent individual motors in experiments. However, the system parameters are based on those of biological systems, see Tables I. The WCA potential is used with the interaction cutoff at 2^1^*/*^6^*σ*, such that the potential between MTs is purely repulsive. The bond stiffness is large, such that the contour length of the MTs remains approximately constant throughout a simulation run. The angle potential is chosen such that MTs are rigid; the persistence length is *𝓁*_*p*_ = 200*L*. We use a time step of duration *δt* = 5.31 *×* 10^-6^*τ*_*R*_. Each run for a particular parameter set consists of 3 10^7^ time steps.

**Table I.** Parameter values used in the simulations.

| Parameter | Symbol | Value | Notes/Biological Values |
| --- | --- | --- | --- |
| Thermal energy | $k_B T$ | 4.11 pN nm | room temperature |
| MT length | $L$ | 0.625 $\mu\text{m}$ | $2.5 \pm 1.4 \mu\text{m}$ [62] |
| MT diameter | $\sigma$ | 25 nm | [63] |
| MT bond angle constant | $\kappa$ | $2.055 \times 10^4 \text{ pN nm}^2$ | rigid MTs |
| MT bond spring constant | $k_s$ | 13.15 pN/nm | preserves MT length [64] |
| Dynamic viscosity | $\eta$ | 1 Pa s | viscosity of cytoplasm [65] |
| Characteristic energy of WCA potential | $\epsilon$ | 4.11 pN nm | [66–68] |
| Motor spring constant | $k_m$ | $6.6 \times 10^{-3} \text{ pN/nm}$ | 0.33 pN/nm per kinesin [69], high number of effective motors |
| Equilibrium motor length | $d_{\text{eq}}$ | 25 nm | MT-MT distance at contact |
| Motor dwell time | $\delta t$ | $4.16 \times 10^{-4} \text{ s}$ | |

We nondimensionalise the key parameters using the MT diameter *σ* or length *L*, thermal energy *k*_B_*T*, and the single-MT rotational diffusion time *τ*_*R*_, see Table II.

**Table II.**
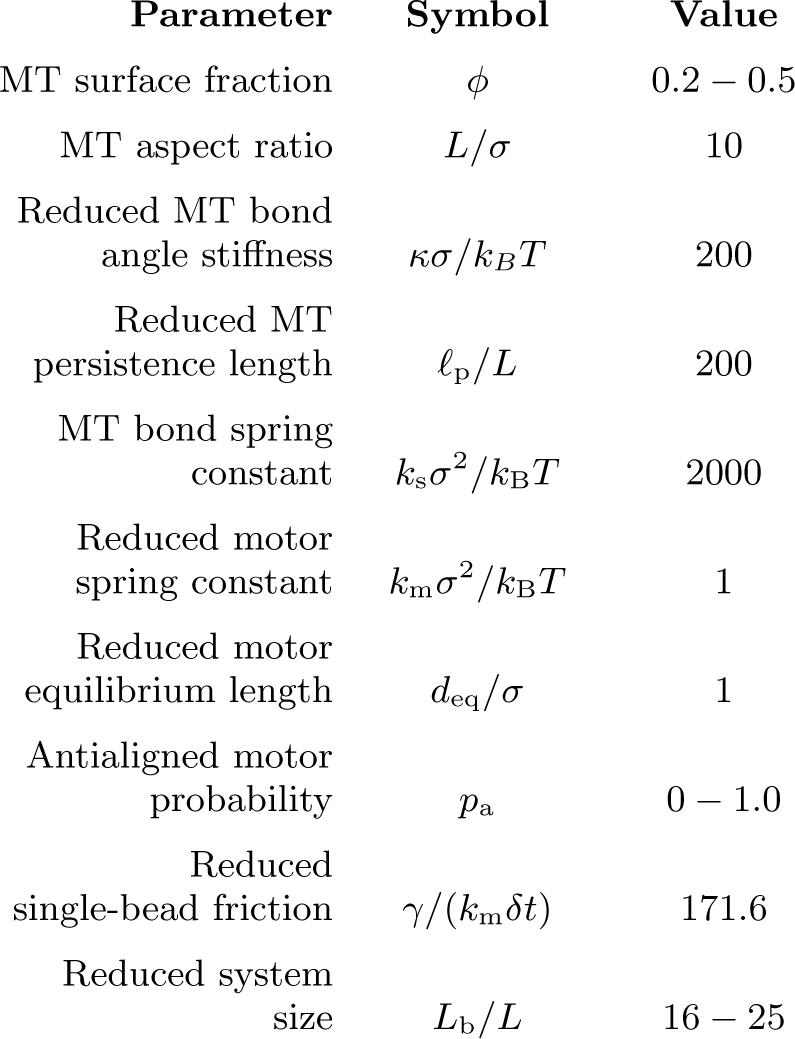
Dimensionless parameters and ranges of the values used in the simulations.

## III. RESULTS

We characterise several distinct processes comprising the phenomenology of active MT motion that arise because of the sliding of adjacent, antialigned MT pairs.

### A. Regimes of MT dynamics

Figure 3 provides an overview of the processes that comprise MT streaming. The fundamental sliding mechanism, which is imposed through the effective motor potential, is the process which occurs at the sliding time *τ*_*N,min*_. The three processes that occur on longer time scales are characterised by the polarity-inversion time *τ*_*Q/*2_, the activity time *τ*^*\**^, and the collective-migration time *τ*_*N,*max_. The active-rotation time *τ*_*r*_ characterises the time when an active MT reaches the end of a polar-ordered domain and changes its orientation.

Figure 3(a) shows a simulation snapshot, which displays multiple, small, polar-aligned MT domains with dynamic interfaces of antialigned MTs between them. The domains are formed by polarity sorting [71] and are in dynamic equilibrium due to MTs that perpetually enter and leave them, see Fig. 3(b). Tracing the individual trajectories shows that MT dynamics consists of a fast streaming mode and a slow diffusion mode, see Fig. 3(c). Uncorrelated displacements in time correspond to slow diffusion within a polar-aligned domain of MTs, and correlated displacements correspond to fast, ballistic streaming at interfaces between domains. This leads to a highly dynamic overall MT structure illustrated by Video 4. At steady state, the length of antialigned interfaces and the size of polar-aligned domains remain constant. The polarity-inversion time *τ*_*Q/*2_ characterises the duration that MTs stay in polar-aligned bundles or in antialigned streams.

Video 3. Video of the trajectory of three selected MTs coloured based on the correlation of adjacent steps in their velocity. The fast-streaming and slow-diffusion modes correspond with the yellow and red parts of the trajectories respectively, according to Fig. 3(c). The video is recorded over a duration of 300*τR*.

Video 4. Simulation video with MTs coloured based on their displacement, where fast displacing MTs are coloured yellow, and slow MTs are coloured blue. The video is recorded over a duration of 100*τR*.

Figure 3(d) shows the MT mean squared displacement

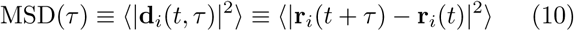

for various motor probabilities [72].

Here, **r**_*i*_(*t*) is the center-of-mass position vector of MT *i* at time *t*, and **d**_*i*_(*t, τ*) is the displacement vector of the center-of-mass of MT *i* between *t* and *t* + *τ*. For passive MTs, *p*_a_ = 0.0, the ballistic regime MSD *∝ τ* ^2^ at short times due to inertia is followed by a diffusive regime MSD *∝ τ* where the MT velocity is dissipated by the environment. For all simulations at finite *p*_a_, we find a superdiffusive regime 10^-1^; ≾ *τ/τ*_*R*_; ≾ 10^1^ with MSD *∝ τ* ^*α*^ and *α >* 1, i.e., with active ballistic motion. Finally, we find a diffusive regime at long times with an active diffusion coefficient that is much higher than for passive Brownian diffusion, see also Sec. III F.

Figure 3(e) shows the histogram of the MT velocity that is projected on the MT orientation vector 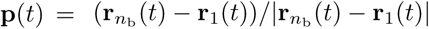,

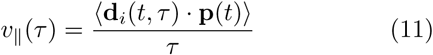

for *p*_a_ = 1.0 and various lag times *τ*. At short lag times, the MT displacement is strongly correlated with the MT’s initial orientation vector and dominated by thermal noise, giving the largest absolute values for *v* _*‖*_ With increasing lag time, the increasing importance of the active motor force is reflected by the increasing asymmetry of the distributions of parallel velocities that are skewed towards positive velocities. At long lag times, the MTs reorient due to active forces, such that both the width of the velocity distribution and the skew again decrease. We characterise the time delay that corresponds to the maximum skew as collective-migration time *τ*_*N,*max_. This time characterises collective motion of neighbouring MTs with similar orientation that travel in the same direction.

Finally, the active orientational correlation time *τ*_*r*_ for MTs is denoted by *τ*_*r*_. This time characterises the crossover between the active-ballistic and the active-diffusive regime in Fig. 3(d). It therefore increases both with increasing size of polar or nematic domains as well as with decreasing rod activity at the interfaces.

### B. Microtubule sliding

The displacement-correlation function,

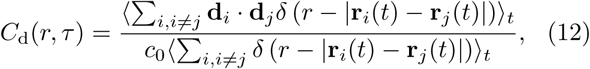

quantifies both spatial and temporal correlations of MT motion. Here, **d**_*i*_ = **r**_*i*_(*t*+*τ*) *-***r**_*i*_(*t*), and 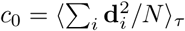 is used for normalisation. Figure 4(a) shows displacement correlation functions for various lag times. At short times and distances, we find negative displacement correlations due to the effective motor potential, which selectively displaces neighbouring antialigned MTs. These negative correlations decay rapidly in space and do not contribute substantially for *r/L* = 1. At intermediate lag times we find positive displacement correlations with a slower spatial decay, and at long lag times no correlations. In the limit *τ →*0, *C*_d_ is the equal-time spatial velocity correlation function [73].

**Figure 4.**
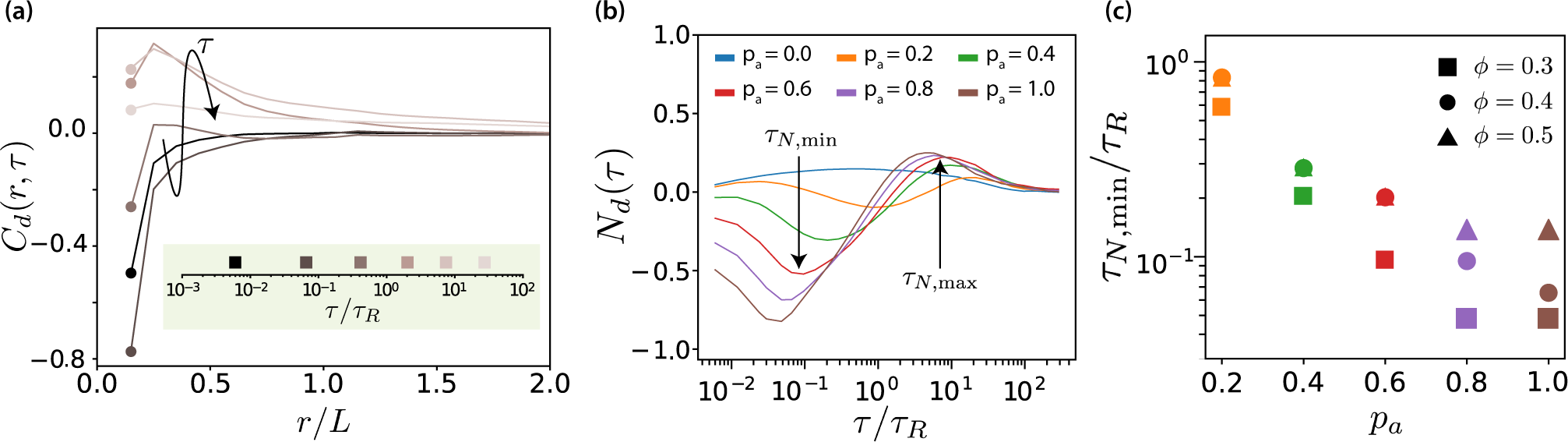
(a) Spatio-temporal correlation function *C*_d_(*r, τ*) for *ϕ* = 0.3 and *p*_a_ = 1.0, for some selected lag times. The arrow and the colours of the curves indicate increasing lag time. The lag times are picked from a logarithmic scale. (b) Neighbour correlation function *N*_d_(*τ*) = *C*_d_(*σ, τ*) for *ϕ* = 0.3 and various *p*_a_ values. (c) The sliding time scale indicated by *τ*_*N,*min_ is shown for various MT surface fractions and *p*_a_ values.

The neighbour displacement correlation function *N*_d_(*τ*) = *C*_d_(*σ, τ*) is defined as the displacement-displacement correlation function at contact *C*_d_(*σ, τ*) [73, 74]. Figure 4(b) shows neighbour displacement correlation functions for various values of *p*_a_ and *ϕ* = 0.3. Firstly, for passive systems, *N*_*d*_ is positive for all MT surface fractions but considerably weaker compared to the correlatiosn in active systems. The small positive correlation is due to steric interaction, and friction due to the roughness of MTs (made up of overlapping beads). For active systems, Fig. 4(b) illustrates that the temporal dependence of *N*_*d*_(*τ*) displays three regimes: for short times, *N*_*d*_(*τ*) is negative and MTs slide antiparallel, for intermediate times, *N*_*d*_(*τ*) is positive and MTs move collectively, and for long times, *N*_*d*_(*τ*) tends to zero and there is no coordinated motion. We focus here on the first regime, whereas the other regimes will be discussed in later sections.

In the short-time regime, *τ/τ*_*R*_ *∼* 10^−1^, the effective motor potential propels neighbouring antialigned MTs away from each other and *N*_d_ is negative. This is aided by higher *p*_a_ but hindered by higher *ϕ*, which opposes active motion sterically. The times *τ*_N,min_ at which the minima occur represent when MTs are propelled because of effective motor interactions, due to presence of the antialigned neighbours. At this time MTs move a small fraction of their length. In Fig. 4(c) the sliding times are collected for different MT surface fractions showing that the sliding is strongly enhanced by activity, where *τ*_N,min_ decreases approximately exponentially with *p*_*a*_ and increases with surface fraction.

### C. Polarity inversion of local environment

In order to characterise an MT’s neighbourhood, we define a pairwise motor partition function [12, 14] which, in the discrete form is

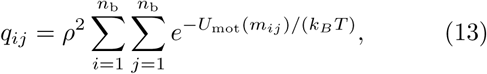

where *ρ* = *n*_b_*/L* is the linear density of binding sites on a single MT, and *m*_*ij*_ is the extension of the motor bound at positions *s*_*i*_ and *s*_*j*_ on different MTs *i* and *j*, respectively [14]. Local polar order weighs pairwise interactions of MTs on the basis of motor binding site availability. It is a function of relative orientation and distance between the beads that are used to model the MTs. Because of the Boltzmann weight, *q*_*ij*_ is significant only for pairs of MTs in close proximity. When two MTs are perfectly overlapping each other, *q*_*ij*_ = 1. When two MTs are sufficiently far away, such that no motors can crosslink them, *q*_*ij*_ = 0, because the MTs are outside motor cutoff range. Since the motor energy *U*_mot_(*m*_*ij*_) increases quadratically with increasing motor extension, the partition function *q*_*ij*_ decays rapidly for increasing distance between the binding sites on the MTs.

The polarity of an MTs environment is quantified by the local polar order parameter *ψ*(*i*). MTs within motor cut-off range are defined to be antialigned if (**p**_*i*_ **p**_*j*_) *<* 0 and polar-aligned if (**p**_*i*_ **p**_*j*_) *≥* 0. By taking the sum of all interacting MTs *j* ≠ *i* with MT *i* [12, 14], we ensure that the local polar order parameter

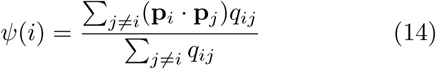

depends on the polarity of the neighbourhood of MT *i*. Here *q*_*ij*_ is given by Eq. (13). The environment of the MT can now be classified into polar (subscript-*P*, 0.5 *< ψ*(*τ*) *<* 1), antipolar (subscript-*A*, *-*1 *< ψ*(*τ*) *< -*0.5), and mixed (subscript-*M*, *-*0.5 *< ψ*(*τ*) *<* 0.5). When a single MT’s environment changes from predominantly antipolar to polar its active motion is stopped and it only moves diffusively.

By tracking changes in *ψ*_*i*_ for single MTs, we measure the time that MTs spend in antialigned or polar-aligned environments, and how this can be affected by *p*_a_ and *ϕ*. The change in local polar order of MT *i* can be written as,

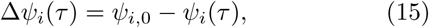

where *ψ*_*i,*0_ = *ψ*_*i*_(*τ* = 0). Figure 5(a) shows ⟨*ψ*_*i,A*_(*τ*) ⟩ and ⟨*ψ*_*i,P*_ (*τ*) ⟩ for *p*_a_ = 1.0 and *p*_a_ = 0.0. In both cases, we find that ⟨*ψ*_*i,A*_(*τ*) ⟩ increases with time, indicating antialigned MTs leaving their antialigned environments, and that ⟨*ψ*_*i,P*_ (*τ*) ⟩ decreases with time, indicating polar-aligned MTs leaving their polar-aligned environments. At long times, *ψ*_*i,A*_ and *ψ*_*i,P*_ converge to the long-time mean ⟨*ψ*_*i,∞*_ ⟩= 0 for passive systems, and to ⟨*ψ*_*i,∞*_(*τ*) ⟩*>* 0 for active systems. The time scale for relaxing ⟨*ψ*_*i*_ ⟩ to the equilibrium value is, as expected, shorter for the active than for the passive system.

**Figure 5.**
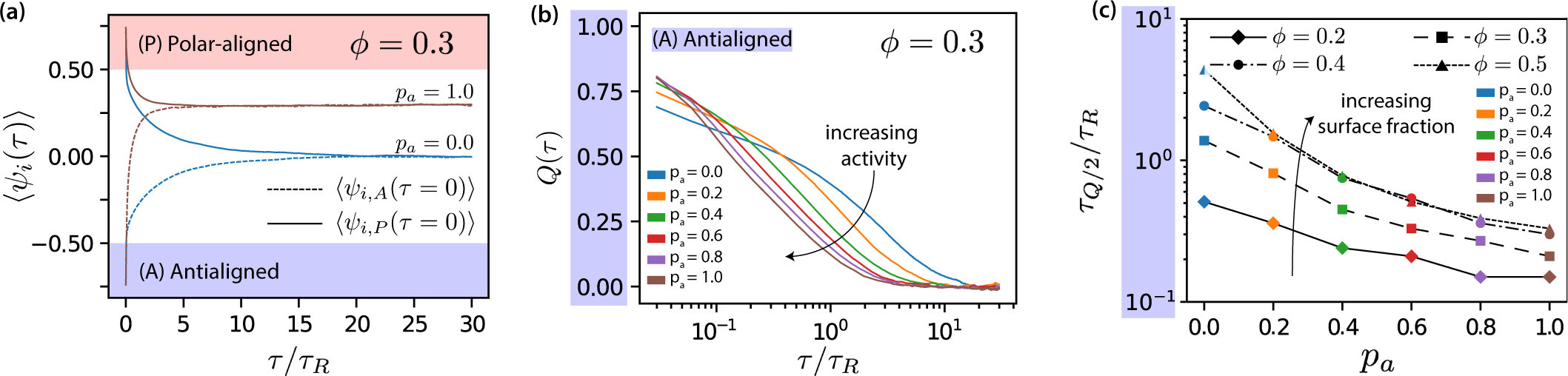
(a) Mean local polar order ⟨*ψi*(*τ*) ⟩ for *p*_a_ = 0.0 and *p*_a_ = 1.0 at *ϕ* = 0.3, for MTs starting from antialigned (dotted line) and aligned (solid line) environments at *τ* = 0. (b) Deviation of local polar order *Q*(*τ*) for *ϕ* = 0.3 for various *p*_a_ for antialigned MTs. (c) Relaxation time for the polar order parameter, *τ*_*Q/*2_ for various *p*_a_ and *ϕ*, estimated by the time for *Q* to decrease to half its initial value.

In order to quantify the change in ⟨*ψ*_*i*_ ⟩, we construct the deviation of local polar order,

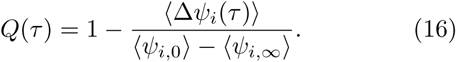

*Q*(*τ*) for the antialigned MTs is shown in Fig. 5(b); the lag time for that *Q*(*τ*) reaches half its initial value is *τ*_*Q/*2*,A*_. While MTs that stay within a polar-ordered domain determine the offset for *ϕ*_*i,A*_ at long times, only MTs entering polar aligned domains determine *τ*_*Q/*2*,A*_. Figure 5(c) shows that *τ*_*Q/*2*,A*_ decreases almost exponentially with *p*_a_ and increases with *ϕ*. In stationary state, the time scales for the inversion of local polar order of initially antialigned and initially polar-aligned MTs are equal, *τ*_*Q/*2*,A*_ = *τ*_*Q/*2*,P*_, compare Fig. 5(b) and Fig. S6. How ⟨*ψ*_*i,∞*_ ⟨ is affected by *ϕ* and *p*_a_ is shown in Fig. S5 in the supplementary material.

### D. Maximal activity

The mean-squared displacements of MTs are ballistic, diffusive, or superdiffusive depending on the lag time, see Fig. 3(d). This is reflected in the distributions of the parallel velocity *v*_‖_, see Eq. (11) and Fig. 3(e). The *v*_‖_ distributions become increasingly asymmetric with increasing lag time when active propulsion dominates over Brownian motion for antialigned MTs–and again less asymmetric when the lag time is further increased and orientational memory is lost. Because of the high number of parallel MTs in our simulations, the position of the main peak is at ⟨*v*_‖_ ⟩= 0 as expected for passive MTs. A skew of the distribution can then be understood as a superposition of a high peak of non-propelled polar-aligned MTs and a small peak that is shifted to positive values of *v*_‖_ for antialigned MTs.

In Fig. 6, we plot skews of *v*_‖_ distributions (Fig. 3(e)) as function of lag time for various *p*_a_ values for *ϕ* = 0.3. The lag time at which the skew of the *v*_‖_ distribution is maximal is defined as the activity time *τ*, where displacements due to active forces are largest relative to displacements due to thermal forces. This activity time *τ*^*\**^ falls into the regime where the MSD is most superdiffusive, see Fig. 3(d). As shown in Fig. 6(b), *τ*^*\**^ exponentially decreases with increasing *p*_a_ and increases with *ϕ*. Increasing motor concentration, and thus a higher amount of active forces in the system, is akin to exponentially shifting the activity time to shorter values.

**Figure 6.**
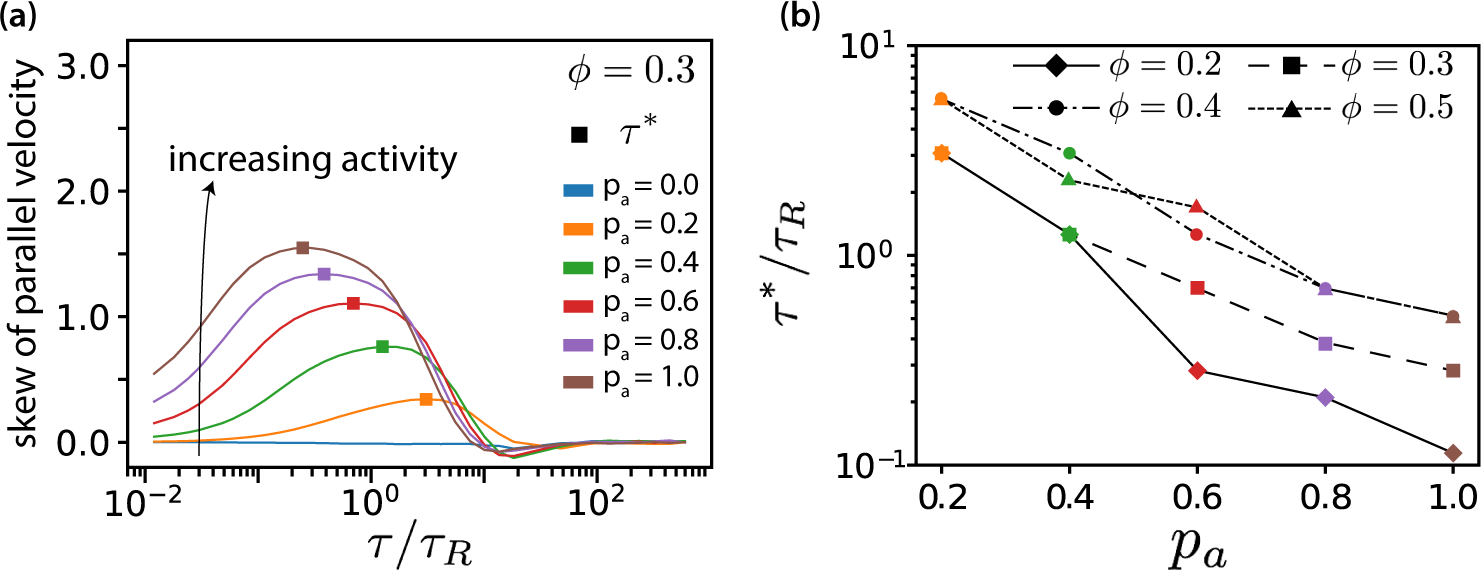
(a) Skew of parallel velocity (*v*_‖_) distribution computed as function of lag times for different *p*_a_ for *ϕ* = 0.3. The probability distributions that correspond to the maximal skew are shown in Fig. S11 together with distributions for few other lag times. (b) Lag time at which maximal skew is observed in the *v*_‖_ (*τ*) distribution (compare Fig. S17). The ordinate is log-scaled to show that *τ*^*\**^ is exponentially decreasing with *p*_a_.

The proportion of aligned (passive) and antialigned (propelled) MTs depends strongly on the area fraction, where the number of antialigned MTs decreases with surface fraction *ϕ*. This corresponds to larger domain sizes and less interfaces with increasing *ϕ*, see Fig. S1 and Fig. S4 in the supplementary material. Further, we notice that for all surface fractions *ϕ* increasing *p*_a_ widens the *v*_‖_ (*τ*^*\**^) distributions, see also Figure S11 in the Supporting Information. The widening of the distribution becomes less pronounced with increasing *ϕ*. The shifting of the negative part of the *v*_‖_ distribution (*v*_‖_ (*τ*^*\**^) *<* 0) to more negative values with increasing *p*_a_ is because *τ*^*\**^ decreases simultaneously.

### E. Collective migration

The observables discussed so far characterise the motion of individual MTs. They only take collective effects into consideration indirectly, *e.g.*, via the asymmetry of the *v*_‖_ distribution for polar-aligned MTs. For a direct discussion of the time scale of collective effects, we return to Fig. 4(b). For times around *τ/τ*_*R*_ = 10, in the intermediate time regime, we observe a positive neighbour displacement correlation. This behaviour is altogether absent at low surface fractions, *ϕ* = 0.2, but for larger *ϕ* the positive correlations increase with increasing *ϕ*. This suggests that neighbouring MTs in a particular stream (likely polar-aligned) travel in the same direction. These polar-aligned MTs will collectively migrate in the same direction because they are in a similarly antialigned environment, i.e., at the same interface with another domain. Correlations in their motion can only manifest at longer lag times, since at short lag times the correlation contribution will be dominated by fast-moving antialigned MTs. We denote the lag time for the maximum of *N*_d_(*τ*), when collective migration occurs, *τ*_N,max_.

In order to explicitly show that positive neighbour displacement correlations observed in the intermediate time regime are due to collective migration of similarly oriented MTs, we can predict the results of photobleaching or photoactivation experiments [12, 75, 76]. Experimentally, in a photobleaching experiment a high-intensity laser beam can be used to inactivate fluorescent molecules in a circular region [77]. The time evolution of the distribution of the light-inactivated regions gives clues about the underlying mechanisms which mediate this motion. Figure 7(a) illustrates that we expect little or no MT sliding to occur in a polar-aligned region, and the photobleached area maintains its shape. In Fig. 7(b), however, the photobleached area is antialigned and we expect bundles of antialigned domains to slide away, causing the photobleached spot to separate into two elongated regions. In our simulations, we perform a similar measurement, where instead of inactivating regions to inhibit fluorescence, we selectively label MT beads within a certain region. We then track their locations for *t* = *τ*_*N,*max_ and investigate their displacements. In Fig. 7(c), for *ϕ* = 0.4, *p*_a_ = 1.0, we visualise a four-MT length radius circular area. MTs move in response to the effective motor potential, and form streams.

**Figure 7.**
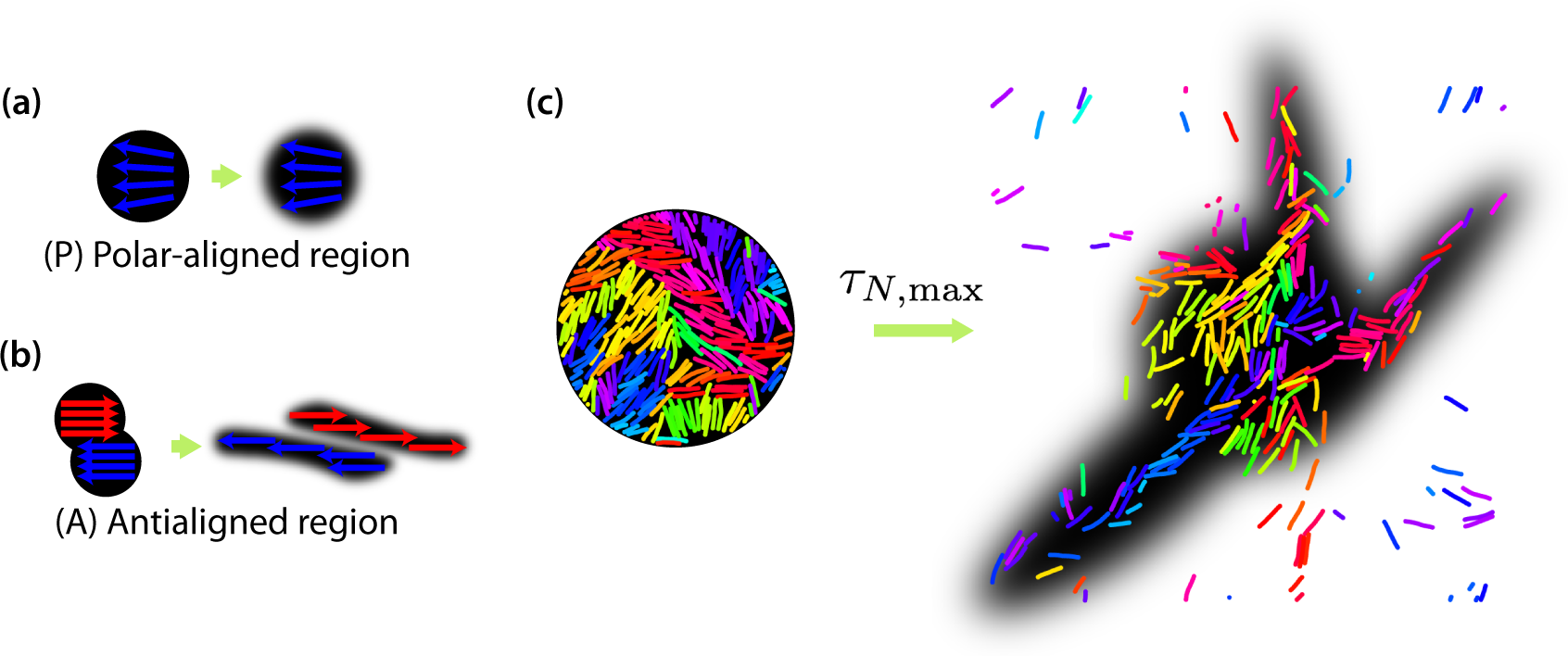
Schematic of expected evolution of photobleached regions in (a) polar-aligned and (b) antialigned regions. (c) Selectively visualised MTs in a circular region within the simulation box, and their evolution after a time of *τN,*_max_, for *ϕ* = 0.4 and *p*_a_ = 1.0. The black backgrounds are predictions of FRAP results.

### F. Active rotation

The longest relevant time scale for the MT dynamics is that of active rotational motion characterised by the orientational correlation function

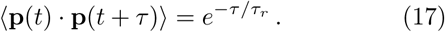

By fitting Eq. (17) to the simulation data (Fig. 8(a)), we obtain the transition time to long-time active diffusive behaviour, *τ*_*r*_. Figure 8(b) shows *τ*_*r*_ for various *p*_*a*_. For passive systems (*p*_*a*_ = 0), *τ*_*r*_ increases with MT surface density *ϕ*. For active systems, *τ*_*r*_ decreases with increasing *p*_*a*_ and with decreasing *ϕ*. In the nematic state for *ϕ* = 0.4 (Fig. S1 in the supplementary material), MTs are no longer able to rotate freely as in the isotropic case.

**Figure 8.**
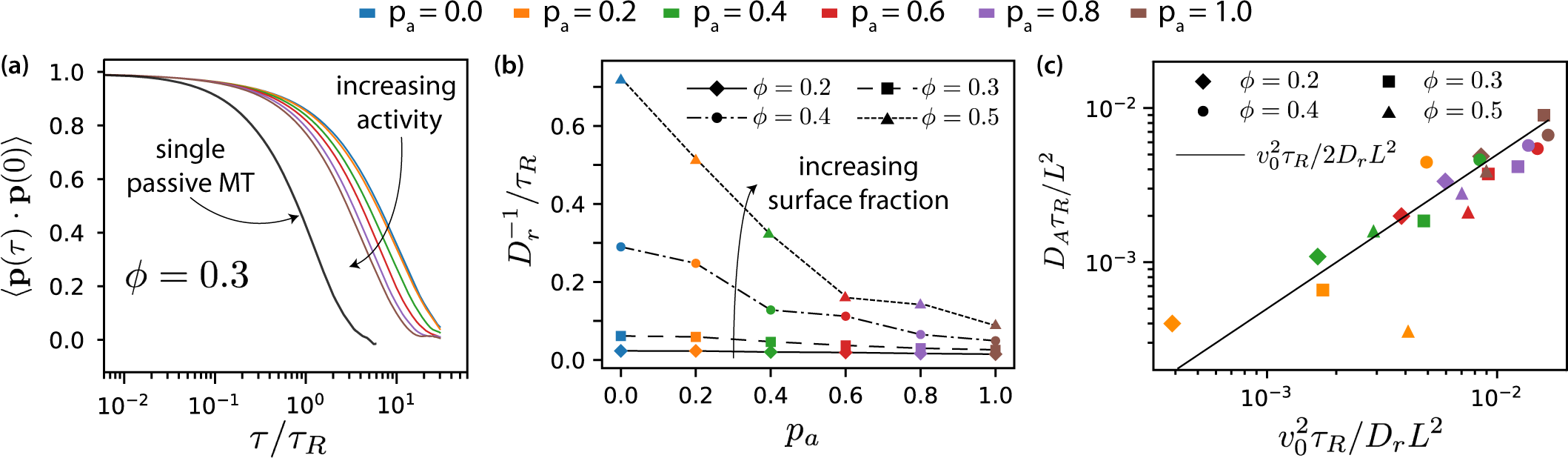
(a) Orientational correlation function for *ϕ* = 0.3 for various *pa*. (b) Inverse of rotational diffusion, *τr* for various *pa* and *ϕ* (c) Active diffusion coefficient *DA* for *p*_a_ = 1.

The decrease of *τ*_*r*_ with increasing *p*_*a*_ is more pronounced at higher MT surface fractions. Smaller values of *τ*_*r*_ correspond to smaller domain sizes. In larger domains, the streams appear at interfaces between polar-ordered domains and antialigned MTs, at larger length scales. The streams extend in the same direction over larger lengths, for longer times, and MTs do not rotate away from their initial orientation as quickly. Also, MTs that are trapped in aligned MT bundles are less likely to exit their environments and their rotational diffusion is smaller for higher *ϕ* and lower *p*_*a*_. For times larger than *τ*_*r*_, MTs diffuse with an active diffusion coefficient 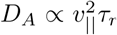 as predicted by the theory of active Brownian particles [78] and is illustrated in Fig. 3(d), see also section V of the supplementary material.

## IV. DISCUSSION AND CONCLUSIONS

In our two-dimensional simulation model, dipolar effective motor forces that drive antialigned MT pairs are sufficient to bring about MT streams which are perpetually created and annihilated, akin to MT streaming in biology. Processes that occur on several characteristic times characterise streaming in our MT-motor mixtures: the characteristic time *τ*_*N,min*_ corresponds to the strongest anti-aligned motion of neighbouring MTs, the time *τ*_*Q/*2_ that an MT stays within a stream, the time *τ*^*\**^ that corresponds to maximal skew of the MT velocity distribution, the collective migration time *τ*_*N,max*_ that characterises maximal directed active motion, and the active rotation time *τ*_r_ that corresponds to single rods traveling the distance of a polar-aligned domain when they loose their orientational memory. Figure 9 and Table III summarise our findings [79].

**Figure 9.**
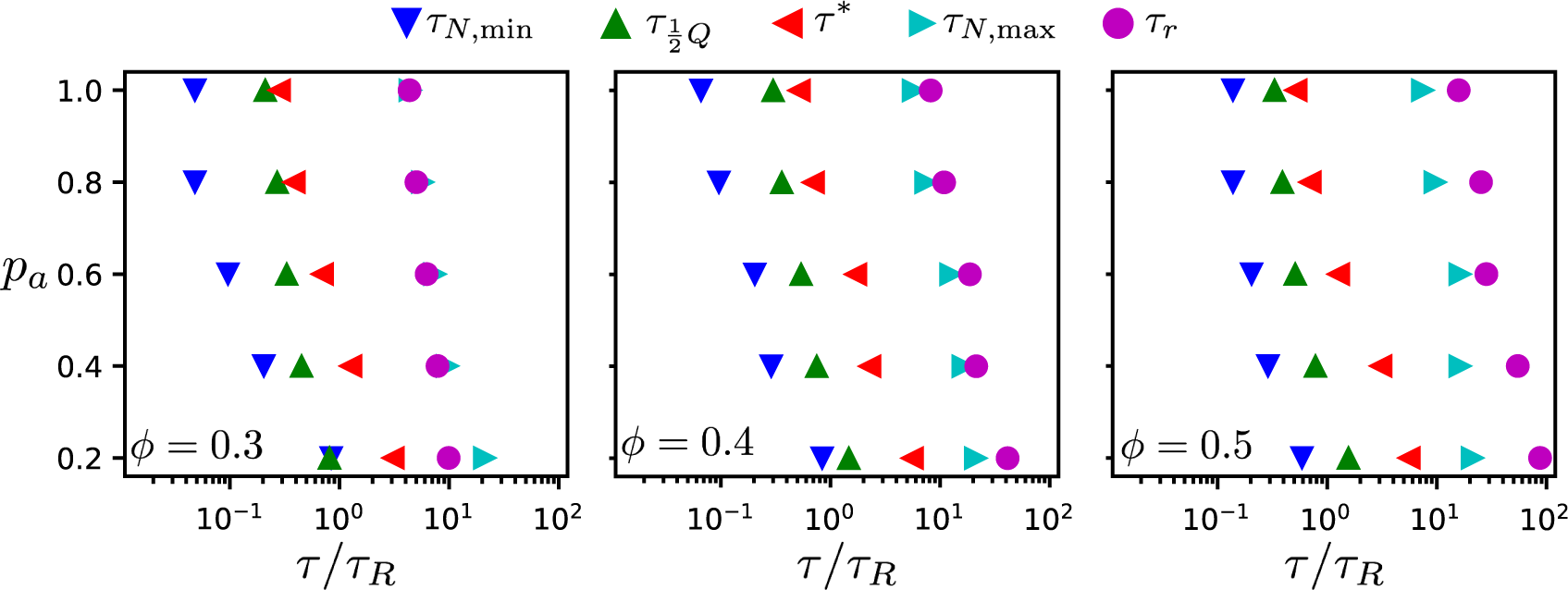
Chronology of events from antialigned MT propulsion to MT rotation (left to right) which make up the streaming process, for various activities *pa* and surface fractions *ϕ* = 0.3, *ϕ* = 0.4, and (c) *ϕ* = 0.5 as indicated.

**Table III.**
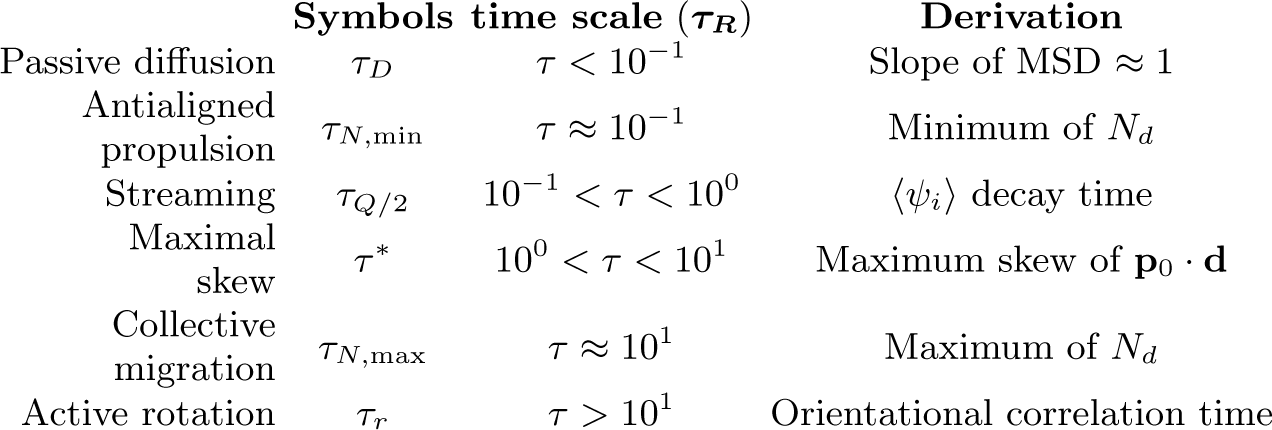
Table of time scales involved in MT dynamics. The time scales reported are approximate values for various *p*_a_ and *ϕ*.

All characteristic times increase with decreasing motor attachment probability *p*_*a*_. We expect the sliding time *τ*_*N,min*_ and the active rotation time *τ*_*r*_ to diverge with vanishing motor attachment probability, while the collective migration time *τ*_*N,max*_ and the polarity-inversion time *τ*_*Q/*2_ attain finite values due to thermal motion and steric interaction between the MTs. The activity time *τ*^*\**^ is not defined for passive systems. Overall, the characteristic times increase with increasing MT surface density, because the lifetime and the coherence of the streams increases. However, our simulations also reveal details of the multi-scale process of streaming in MT-motor mixtures. For example, the time for an MT to transition from a polar-aligned to an antialigned environment is similar to the sliding time for low motor forces, where thermal motion dominates, and to the activity time for high motor forces when the streams are more stable.

The closest ‘bottom-up’ experimental system to our simulation model is the *in vitro* model system of microtubule bundles, kinesin complexes, and depletants at the oil-water interface, investigated in Ref. [22]. A detailed quantitative comparison is currently not possible because the characteristic length scales in simulations and experiment are quite different. The depletion-induced MT bundle formation in the experiments leads to a characteristic length scale of the order of 10*µm*, whereas the MTs in the simulations have lengths below 1*µm*. However, on a more qualitative level interesting correspondences are revealed. The transition between diffusive and ballistic MSDs in the simulations has also been reported for the *in vitro* model system [22]. This allows the comparison of the active diffusive regime for times *τ > τ*_*r*_ and for lengths longer than a typical domain size. Whereas the motion in suspensions of passive MTs is diffusive, a ballistic regime at large lag times is found for increasing concentrations of active motors (simulations) and for increasing ATP concentration (experiment), see Supplementary Information.

Some of our results can be used to interpret experimental results *in vivo*. For example, using Particle Image Velocimetry (PIV) in *Drosophila* cells, fluid velocity distributions have been measured for wild-type oocytes and those lacking *pat1*, a protein required for kinesin heavy chain to maximise its motility [34]. The main peak is close to a velocity of 10 nm/s, which hints that the majority of the MTs are propelled. As in our simulations, heavy tails in the velocity distribution have been reported in the experiments. Comparing the experimental data for wild-type and *pat1* -deficient systems, the mean speed of the wild-type system was slower and the velocity distribution had heavier tails. This qualitatively agrees with our findings for varying *p*_a_. It was suggested in Ref. [34] that the heavy tails in the velocity distribution of the cytosol reflect a combination of an underlying distribution of motor speeds, and a complex MT network geometry. From our simulations, we conclude that neither a complex three-dimensional cytoskeletal geometry nor a combination of different motor speeds are required to reproduce cytoskeletal velocity distributions with heavy tails.

We have studied the characteristic times of MT-motor dynamics relevant for the cytoskeleton using a coarsegrained motor model and Langevin Dynamics simulations in the overdamped regime. This allows us to access both the single-MT level as well as the collective-MT level. In previous studies that use a similar coarsegraining technique for the motor activity, the focus has been on understanding and capturing biologically relevant cytoskeletal structures [42, 50]. Here, for the first time, we have decomposed the time scales of activity from single MTs to system-scale ordering and streaming.

MT advection has also been analysed using photoconversion in interphase *Drosophila* S2 cells, where MTs were observed to buckle and loop [9]. MT motion was visualised by photoconverting a circular region within the cell. These MTs were observed over a 7 minute period, during which 36% of the MTs were determined to be motile. It was observed that MTs spent most of the time not moving, but underwent abrupt long-distance streaming. They were found to achieve velocities up to 13 *µ*m/min, during these bursts of active motion. These observations are very similar to those in our simulations, where MTs spend most of their times in stable polar-aligned bundles, but when in contact with an antialigned MTs coherently stream over large distances. We find similar fractions of motile MTs between 30% and 40% also for *ϕ* = 0.3 in our simulations. Our study provides the basis for a more detailed quantitative comparison with experiments because the model can be easily extended to include further relevant aspects, such as a 3D cytoskeletal network, crosslinking proteins, and cellular confinement.

Our simulations show collective migration of MTs that is maximal at *τ*_*N,*max_. Using a FRAP-like visualisation of our data, we find elongated MT stream patterns similar to those observed in experiments [9]. This confirms that similarly oriented MTs move colletively in the same stream. Based on the polarity-sorting mechanism of MTs, qualitatively similar FRAP results have previously been predicted using computer simulations [12]. Experimental studies of a system on various length and time scales should allow testing the chronology that we predict. For example, systems with different fractions of fluorescent MTs with fixed lengths should give access to both collective as well as single-MT dynamics, e.g., using FRAP/photoactivation for systems with many labeled MTs to quantify collective dynamics and confocal microscopy for systems with few labeled MTs to investigate correlations in single-MT motion.

We have studied collective motion in active gels based on single MTs. Our spatio-temporal displacement correlation functions show that antialigned MTs slide away from each other in opposite directions for short time windows, while in agreement with experiments positive correlations occur for long time windows [34]. Our twodimensional simulations resemble systems close to an interface that have been used to experimentally study hierarchically assembled active matter [22]. They also lay the foundations for future studies of 3D systems and have allowed us to test parameter regimes using less computationally expensive, two-dimensional systems. Furthermore, although we observe streaming without hydrodynamics, hydrodynamic interactions may still be an important player for motor-MT systems, which can be investigated in future studies.

To summarize, our results provide a direct handle to fully characterise MT streaming over a wide range of time and length scales. Future experimental studies using modern microscopy techniques may allow testing our predictions. Future theoretical and computer simulation studies may provide further insights, such as the importance of the aspect ratio of the MTs, the presence of motors between polar-aligned MTs, and the effect of crosslinkers important for buckling and looping of flexible MTs.

## ACKNOWLEDGMENTS

O. D. and G. S. acknowledge support by the International Helmholtz Research School of Biophysics and Soft Matter (IHRS BioSoft). CPU time allowance from the Jülich Supercomputing Centre (JSC) is gratefully acknowledged.

## Reference

[1] M. E. Quinlan, Annu. Rev. Cell Dev. Biol. 32, 173 (2016).

[2] I. M. Palacios and D. St Johnston, Development 129, 5473 (2002).

[3] W. Lu, M. Winding, M. Lakonishok, J. Wildonger, and I. I. Gelfand, Proc. Natl. Acad. Sci. U.S.A. 113, E4995 (2016).

[4] G. Berthold, Studien über Protoplasmamechanik (Verlag von Arthur Felix, 1886).

[5] H. Gutzeit, Rouxs Arch Dev Biol 195, 173 (1986).

[6] L. R. Serbus, B.-J. Cha, W. E. Theurkauf, and W. M. Saxton, Development 132, 3743 (2005).

[7] C. E. Monteith, M. E. Brunner, I. Djagaeva, A. M. Bielecki, J. M. Deutsch, and W. M. Saxton, Biophys. J. 110, 2053 (2016).

[8] W. E. Theurkauf, S. Smiley, M. Wong, and B. M. Alberts, Development 115, 923 (1992).

[9] A. L. Jolly, H. Kim, D. Srinivasan, M. Lakonishok, A. G. Larson, and V. I. Gelfand, PNAS 107, 12151 (2010).

[10] M. Winding, M. T. Kelliher, W. Lu, J. Wildonger, and V. I. Gelfand, PNAS 113, E4985 (2016).

[11] R. D. Vale and R. A. Milligan, Science 288, 88 (2000).

[12] T. Gao, R. Blackwell, M. A. Glaser, M. D. Betterton, and M. J. Shelley, Phys. Rev. Lett. 114, 048101 (2015).

[13] R. Blackwell, O. Sweezy-Schindler, C. Baldwin, L. E. Hough, M. A. Glaser, and M. D. Betterton, Soft Matter 12, 2676 (2016).

[14] A. Ravichandran, G. A. Vliegenthart, G. Saggiorato, T. Auth, and G. Gompper, Biophys. J. 113, 1121 (2017).

[15] J. M. Belmonte, M. Leptin, and F. Nédélec, Mol. Syst. Biol. 13, 941 (2017).

[16] Ö. Duman, R. E. Isele-Holder, J. Elgeti, and G. Gompper, Soft Matter 14, 4483 (2018).

[17] D. Needleman and Z. Dogic, Nature Reviews Materials 2, 17048 (2017).

[18] A. Zöttl and H. Stark, J. Phys.: Condens. Matter 28, 253001 (2016).

[19] C. Bechinger, R. Di Leonardo, H. Lowen, C. Reichhardt, G. Volpe, and G. Volpe, Rev. Mod. Phys. 88, 045006 (2016).

[20] J. Elgeti, R. G. Winkler, and G. Gompper, Rep. Prog. Phys. 78, 056601 (2015).

[21] K. Chen, B. Wang, and S. Granick, Nat. Mater. 14, 589 EP (2015).

[22] T. Sanchez, D. T. Chen, S. J. DeCamp, M. Heymann, and Z. Dogic, Nature 491, 431 (2012).

[23] S. J. Decamp, G. S. Redner, A. Baskaran, M. F. Hagan, and Z. Dogic, Nature Materials 14, 1110 (2015).

[24] F. G. Woodhouse and R. E. Goldstein, Proc. Natl. Acad. Sci. U.S.A. 110, 14132 (2013).

[25] P. K. Trong, H. Doerflinger, J. Dunkel, D. S. Johnston, and R. E. Goldstein, eLife 4, e06088 (2015).

[26] M. J. Schnitzer and S. M. Block, Nature 388, 386 (1997).

[27] D. Mizuno, C. Tardin, C. F. Schmidt, and F. C. MacKintosh, Science 315, 370 (2007).

[28] D. A. Head, W. Briels, and G. Gompper, BMC Biophys. 4, 18 (2011).

[29] T. Hiraiwa and G. Salbreux, Phys. Rev. Lett. 116, 188101 (2016).

[30] P. Ronceray, C. P. Broedersz, and M. Lenz, Proc. Natl. Acad. Sci. U.S.A. 113, 2827 (2016).

[31] D. A. Head, W. J. Briels, and G. Gompper, Phys. Rev. E 3, 032705 (2014).

[32] S. L. Freedman, S. Banerjee, G. M. Hocky, and A. R. Dinner, Biophys. J. 113, 448 (2017).

[33] E. Toprak, A. Yildiz, M. T. Hoffman, S. S. Rosenfeld, and P. R. Selvin, Proc. Natl. Acad. Sci. U.S.A. 106, 12717 (2009).

[34] S. Ganguly, L. S. Williams, I. M. Palacios, and R. E. Goldstein, PNAS 109, 15109 (2012).

[35] F. Nedelec and D. Foethke, New J. Phys. 9, 427 (2007).

[36] M. Burdyniuk, A. Callegari, M. Mori, F. Nédélec, and P. Lénárt, J. Cell Biol., jcb (2018).

[37] B. Lacroix, G. Letort, L. Pitayu, J. Sallé, M. Stefanutti, G. Maton, A.-M. Ladouceur, J. C. Canman, P. S. Maddox, A. S. Maddox, et al., Dev. Cell 45, 496 (2018).

[38] G. Letort, F. Nedelec, L. Blanchoin, and M. Thery, Mol. Biol. Cell 27, 2833 (2016).

[39] E. Nazockdast, A. Rahimian, D. Needleman, and M. Shelley, Mol. Biol. Cell 28, 3261 (2017).

[40] E. Nazockdast, A. Rahimian, D. Zorin, and M. Shelley, J. Comput. Phys. 329, 173 (2017).

[41] S. Swaminathan, F. Ziebert, I. Aranson, and D. Karpeev, EPL 90, 28001 (2010).

[42] Z. Jia, D. Karpeev, I. S. Aranson, and P. W. Bates, Phys. Rev. E 77, 051905 (2008).

[43] R. G. Winkler and G. Gompper, Handbook of Materials Modeling: Methods: Theory and Modeling, 1 (2018).

[44] K. Müller, D. A. Fedosov, and G. Gompper, J. Comput. Phys. 281, 301 (2015).

[45] G. Gompper, T. Ihle, D. Kroll, and R. Winkler, in Advanced computer simulation approaches for soft matter sciences III (Springer, 2009) pp. 1–87.

[46] R. E. Isele-Holder, J. Jäger, G. Saggiorato, J. Elgeti, and G. Gompper, Soft Matter 12, 8495 (2016).

[47] J. D. Weeks, D. Chandler, and H. C. Andersen, J. Chem. Phys. 54, 5237 (1971).

[48] K. Kruse and F. Jülicher, Phys. Rev. Lett. 85, 1778 (2000).

[49] K. Kruse, J.-F. Joanny, F. Jülicher, J. Prost, and K. Sekimoto, Eur. Phys. J. E 16, 5 (2005).

[50] I. S. Aranson and L. S. Tsimring, Phys. Rev. E 71, 050901 (2005).

[51] I. S. Aranson and L. S. Tsimring, Phys. Rev. E 74, 031915 (2006).

[52] G. Salbreux, J. Prost, and J.-F. Joanny, Phys. Rev. Lett. 103, 058102 (2009).

[53] A. Cördoba, J. D. Schieber, and T. Indei, Soft Matter 11, 38 (2015).

[54] K. Kruse, J. F. Joanny, F. Jülicher, J. Prost, and K. Sekimoto, Physical Review Letters 92, 078101 (2004).

[55] K. Kruse, S. Camalet, and F. Jülicher, Phys. Rev. Lett. 87, 138101 (2001).

[56] Although the force for each motor lasts for a single time step, this duration is not a characteristic time for the motor. Our model describes continuum propulsion forces on MTs imposed by motors. This is supported by Fig. S10 in the supporting information, which shows that the MT parallel velocity is proportional to the fraction of time that motors are active, although the average time that a force is exerted on a particular monomer increases with the fraction of active motors.

[57] F. Jülicher, A. Ajdari, and J. Prost, Rev. Mod. Phys. 69, 1269 (1997).

[58] Similarly, we can also implement motors between polaraligned MTs.

[59] J. Kerssemakers, J. Howard, H. Hess, and S. Diez, Proc. Natl. Acad. Sci. U.S.A. 103, 15812 (2006).

[60] L. Winters, I. Ban, M. Prelogovic, N. Pavin, and I. M. Tolic, bioRxiv (2018), doi:10.1101/347831.

[61] S. Plimpton, J. Comput. Phys. 117, 1 (1995).

[62] J. Howard, A. J. Hudspeth, and R. D. Vale, Nature 342, 154 (1989).

[63] D. Chrétien and R. H. Wade, Biol. Cell 71, 161 (1991).

[64] R. E. Isele-Holder, J. Elgeti, and G. Gompper, Soft Matter 11, 7181 (2015).

[65] D. Wirtz, Annu. Rev. Biophys 38, 301 (2009).

[66] M. A. Bates and D. Frenkel, J. Chem. Phys. 112, 10034 (2000).

[67] P. Bolhuis and D. Frenkel, J. Chem. Phys. 106, 666 (1997).

[68] S. C. McGrother, D. C. Williamson, and G. Jackson, J. Chem. Phys. 104, 6755 (1996).

[69] C. M. Coppin, J. T. Finer, J. A. Spudich, and R. D. Vale, Biophys. J. 68, 242S (1995).

[70] M. Abkenar, K. Marx, T. Auth, and G. Gompper, Phys. Rev. E 88, 062314 (2013).

[71] V. Wollrab, J. M. Belmonte, M. Leptin, F. Nédélec, and G. H. Koenderink, bioRxiv, 314484 (2018).

[72] We use m = 1 and γ = 1 (in simulation units). The single MT center-of-mass dynamics is overdamped for times t/τR > 2m/γτR 10−2, which is shorter than the shortest time scale of interest shown in Fig. 3(d).

[73] A. Wysocki, R. G. Winkler, and G. Gompper, Europhys. Lett. 105, 48004 (2014).

[74] B. Doliwa and A. Heuer, Phys. Rev. E 61, 6898 (2000).

[75] T. Mitchison, J. Cell Biol. 109, 637 (1989).

[76] J. Hush, P. Wadsworth, D. Callaham, and P. Hepler, J. Cell Sci. 107, 775 (1994).

[77] D. Axelrod, D. Koppel, J. Schlessinger, E. Elson, and W. W. Webb, Biophys. J. 16, 1055 (1976).

[78] J. R. Howse, R. A. L. Jones, A. J. Ryan, T. Gough, R. Vafabakhsh, and R. Golestanian, Phys. Rev. Lett. 99, 048102 (2007).

[79] We do not show φ = 0.2, because there is no evidence of streaming in these systems, and the chronology of events are not consistent with those observed for higher surface fractions.

