## Supplemental Information for "Chronology of motor-mediated microtubule streaming"

### SUPPLEMENTARY MATERIAL

A. Ravichandran,[1] Ö. Duman,[1] M. Hoore,[1] G. Saggiorato,[1, 2] G. A. Vliegenthart,[1] T. Auth,[1] and G. Gompper[1]

[1] *Theoretical Soft Matter and Biophysics, Institute of Complex Systems and Institute for Advanced Simulation, Forschungszentrum Jülich, 52425 Jülich, Germany*

[2] *LPTMS, CNRS, Univ. Paris-Sud, Université Paris-Saclay, 91405 Orsay, France*

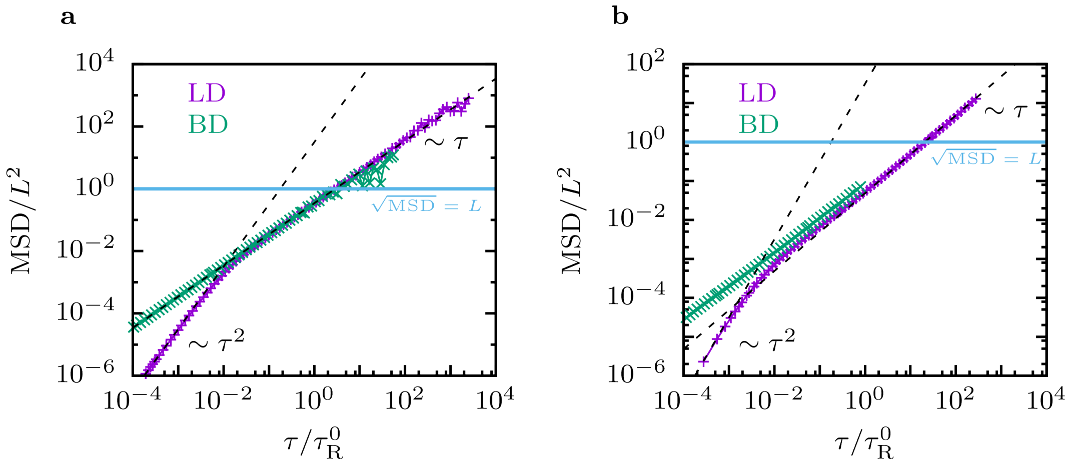


FIG. S1. **a** MSD of single filament calculated with Langevin dynamics and with Brownian dynamics. **b** MSD of a concentrated filament suspension ($\phi =$) calculated with Langevin dynamics and with Brownian dynamics.

### I. LANGEVIN DYNAMICS VERSUS BROWNIAN DYNAMICS

For a single stiff filament made of $n_\mathrm{b}$ beads with mass $m$ the center-of-mass mean squared displacement in the inertial regime is $\mathrm{MSD} = 2(k_\mathrm{B}T/n_\mathrm{b}m)t^2$ and in the diffusive regime $\mathrm{MSD} = 4(k_\mathrm{B}T/n_\mathrm{b}\gamma)t$ such that the crossover time from ballistic to diffusive motion is $t_\mathrm{co} = 2m/\gamma$. The single filament (passive) rotation time is obtained from the orientational correlation function which in the diffusive regime for a rigid rod made of $n_b$ beads decays like $C(t) = \exp(-t/\tau_\mathrm{R})$ with $\tau_\mathrm{R} = 2\gamma r_0^2 f(n_b)/k_BT$ and $f(n_b) = n_b(n_b^2 - 1)/24$. For the crossover time from ballistic to diffusive we find then $t_\mathrm{co}/\tau_\mathrm{R} \approx 0.01$, see Fig. S1**a**. For concentrated systems the diffusive part of the MSD will shift downwards (to a smaller diffusion coefficient) and $t_\mathrm{co}/\tau_\mathrm{R}$ will shift to shorter times, see Fig. S1**b**. All relevant time scales discussed in the main text are larger than $\tau/\tau_R = 0.01$ and therefore, the inertial term in the Langevin equation is not relevant for the processes that we have studied.

### II. ORIENTATIONAL ORDER AND DOMAIN FORMATION: SIMULATION SNAPSHOTS

In this study, MT sliding is driven by an effective motor potential. The qualitative behaviour of our systems is similar to results from an explicit motor model [1]. In both cases, a perfectly sorted state is not achieved, which leads to persistent motion. Both, motor activity and steric interactions determine the dynamics of the systems, see Fig. S2. Activity increases with MT surface fraction at small surface fractions if a lack of motors between nearby antialigned MTs limits active dynamics. However, at high surface fractions steric effects impede MT motion, and increasing MT surface fraction decreases MT dynamics.

In general, the motor actions are controlled using an antialigned motor probability $p_\mathrm{a}$ and a polar-aligned motor probability $p_\mathrm{p}$. The parameters control whether a motor appears between two MTs beads that have a distance within the cutoff radius for the interaction The antialigned motor probability directly affects MT advection, while the aligned motor probability leads to cohesion of polar-aligned regions. Structures formed purely due to interactions between antialigned MTs are shown in Fig. S2. In cases without motor interactions, $p_\mathrm{a} = 0.0$, the MTs are isotropic at $\phi = 0.3$, and appear to be nematically aligned at $\phi = 0.5$. This is consistent with the phasediagram for hard rods in two dimensions calculated by Bates and Frenkel [2] who estimated the I-N transition for $L/D = 10$ around $\phi \approx 0.5$.

At $\phi = 0.3$, a finite effective motor potential leads to the formation of small and highly dynamic polar-aligned domains. There is no appreciable difference in the systems' structures due to increasing antialigned motor probability. MTs enter and leave the polar-aligned domains continuously because of their high probability to interact with antialigned MTs. At $\phi = 0.4$ and $\phi = 0.5$, MT streaming thus breaks apart the large nematic domains that are observed in the passive system. The size of polar-aligned domains are increased compared to $\phi = 0.3$. For a specific surface fraction, the domain sizes decrease with increasing $p_\mathrm{a}$. We find domains that span the entire simulation box at $0.2 \leq p_\mathrm{a} \leq 0.6$ and $\phi = 0.5$.

### III. QUANTIFICATION OF STRUCTURE AND DYNAMICS

#### A. Local polar order

Calculating the local polar order parameter $\psi_i$ of a MT, defined in Eq. (14) of the main text, helps us to quantify the extent of bundling as function of antialigned motor probability $p_\mathrm{a}$ and MT surface fraction $\phi$. Also, the local polar order parameter correlates with the MT velocity. As shown in Fig. S3, all MTs in the system are separated into three groups based on their local environment:

1. A fraction $n_A$ of MTs is in an antialigned environment with $\psi_i \leq -0.5$.

2. A fraction $n_M$ of MTs is in a mixed environments with $-0.5 < \psi_i \leq 0.5$ [3].

3. A fraction $n_P$ of MTs is in a polar-aligned environment with $0.5 < \psi_i$.

Figure S3 gives examples of MTs in each environment category. Figure S4 quantifies the polar structure of the systems shown in Fig. S2. At the lowest density, $\phi = 0.2$, the largest population of MTs are in a mixed environment. This population of MTs lead to the peak at $\psi_i = 0.0$, whose height decreases with increasing MT surface fraction.

With increasing MT surface fraction, the fraction of MTs that are either antialigned or polar aligned increases. For finite motor probability, due to polarity sorting the number of polar-aligned MTs exceeds the number of antialigned MTs. At $\phi = 0.4$ and $\phi = 0.5$, the majority of all MTs is in a polar environment.

In Fig. S5, we quantify the ratios of antialigned to polar-aligned MTs, and of mixed MTs. For passive systems, $p_\mathrm{a} = 0$, we find $n_A/n_P = 1$ for all surface fractions shown in Fig. S5(a). For systems with antialigned MT sliding, $n_A/n_P$ decreases with increasing surface fraction for all $p_\mathrm{a}$ values. This corresponds to an increase of polar-aligned clusters with increasing MT surface fraction, see Fig. S2. Furthermore, higher $p_\mathrm{a}$ values lead to larger values of $n_A/n_P$. At high surface fractions, the dependence of $n_A/n_P$ on $p_\mathrm{a}$ decreases. However, Fig. S2 shows that the size of polar-aligned domains decreases with activity at $\phi = 0.5$. The ratio $n_A/n_P$ is a good indicator for the length of antialigned MT interface in the system, which corresponds to smaller domain sizes of polar-aligned regions. The decrease in $n_m$, for the passive system in Fig. S5(b) is due to the isotropic-nematic phase transition. In general, we see a similar, but stronger decrease in $n_m$ also for all active systems. At long times, the mean local order parameter increases strongly with MT surface fraction $\phi$ and weakly with the antialigned motor probability $p_\mathrm{a}$, see Fig. S6.

#### B. Mean squared displacements of filaments

We quantify the velocities of MTs for various lag times $\tau$ using their MSDs defined in Eq. (10) of the main text. Figure S8 shows MSD/$\tau$ for various values of $p_\mathrm{a}$ and $\phi$. In all active cases, $p_\mathrm{a} > 0$, there are four different regimes of motion, contrary to only two regimes in the case of a single, passive MT: ballistic motion $\propto \tau^2$ due to inertial effects for short times and diffusive motion $\propto \tau$ at long times. The two additional regimes are an active ballistic regime $\propto \tau^2$ at intermediate times, and an active diffusive regime $\propto \tau$ at long times.

The MT mean squared displacements for the effective-motor system can be qualitatively described using the MSD that has been developed for active Brownian particles [4–6],

$$\langle |\mathbf{d}_i(\tau)|^2 \rangle = 4D\tau + \frac{2v_0^2}{D_r^2}\left(D_r\tau + e^{-D_r\tau} - 1\right), \tag{1}$$

where $D$ is the passive translational diffusion coefficient, $D_r$ is the active rotational diffusion coefficient, $v_0$ is an active velocity of the active particle. (In our case $v_0 = v_{||}$.) At long times,

$$\langle |\mathbf{d}_i(\tau)|^2 \rangle \approx 4D\tau + \frac{2v_0^2}{D_r}\tau \equiv 4D_A\tau. \tag{2}$$

If the passive diffusion coefficient $D$ is much smaller than $v_0^2/D_r$, the active diffusion coefficient $D_A = v_0^2/2D_r$ is solely determined by the active velocity of the MT and the active rotational diffusion coefficient. However, this relationship is effective, because contrary to active Brownian sphere-like or disk-like particles $D_\mathrm{r}$ is not thermal, but instead depends on the active velocity of a filament and the domain structure.

In Fig. [**?** ] we compare the MSDs for micrometer-sized test particles from Ref. [7] with our data for single MTs. Both passive systems show hindered diffusion because of the high MT densities. Increasing the effective motor probability in the simulations corresponds to an increasing ATP concentration in the experiments. At small activities the enhanced diffusion is mostly present at long lag times, whereas for high activities the transition to the active regime shifts to shorter times and the ballistic and–in the simulations–also the active-diffusive regime are observed. MT bundling in the experiments shifts both length and time for the crossover to the active diffusive regime to larger values, such that the active-diffusive regime is not accessible in the experiments.

#### C. Neighbour displacement correlation function

The neighbour displacement correlation function $N_{\mathrm{d}}(\tau)$, discussed in Sec. III B of the main text, describes how initially neighbouring filaments move. Figure S10 shows neighbour displacement functions for several MT surface fractions. For short times, the displacements are anticorrelated for all surface fractions and all non-zero antiparallel motor probabilities. The lag time for the strongest anticorrelation, $\tau_{N,min}$, decreases with $p_{\mathrm{a}}$. The strength of the anticorrelation decreases with increasing $\phi$ and decreasing $p_{\mathrm{a}}$, due to increased steric hindrance and a weaker driving force, respectively. The lag times for the strongest correlation, $\tau_{N,max}$, characterises collective migration of filaments. For the smallest MT surface fraction $\phi = 0.2$ collective migration does not take place, for higher MT surface fractions collective migration increases with increasing steric hindrance.

#### D. Parallel velocity

Figure S11 shows that the parallel velocity $v_{\|}$ of a MT, defined in Eq. (11) of the main text, decreases with increasing duration $\tau$ of the time window for that the velocity is measured. For small MT surface fractions, the time scale on that $v_{\|}$ decreases is the rotational diffusion time for a single MT. For higher MT surface fractions, the orientation of the MTs is more persistent, such that the decrease of $v_{\|}$ with the duration of the time window decreases. Furthermore, while at higher MT surface fractions more effective motors are present in a system, also the hindrance of MT motion by steric interactions increases. In addition, the availability of antialigned MT pairs controls the activity, which depends on the structure of the MT system. The parallel velocity is highest for intermediate MT surface fraction $\phi = 0.3$, see also Fig. S12.

Figure S12 shows that the parallel velocities for $\tau \to 0$ are proportional to the antialigned motor probability $p_{\mathrm{a}}$. This provides a good control over the activity in the system. The results seem to suggest that systems with small MT surface fractions, $\phi = 0.2$ and $\phi = 0.3$, are more active compared with those with high MT surface fractions, $\phi = 0.4$ and $\phi = 0.5$. However, at higher surface fractions the MT deviate slower from their initial orientation, see Fig. S11, which suggests that MTs are more likely to stream.

#### E. Maximal activity

Figure S13 shows histograms of MT parallel velocities for various durations of the time window for the velocity measurements. The distributions are asymmetric, those with the highest skew that correspond to a time window of optimal duration, the activity time $\tau^*$, are marked. Figure S14 shows the parallel-velocity distributions with maximal skews for various antialigned motor probabilities $p_{\mathrm{a}}$ and MT surface fractions $\phi$. The widths of the distributions increase with increasing $p_{\mathrm{a}}$ and decreasing $\phi$. Finally, Fig. S15 shows that the parallel velocities for a time window duration of the activity time $t^*$–unlike the parallel velocities for $\tau \to 0$ shown in Fig. S12–do not strongly depend on the MT surface fraction.

#### F. Fluorescence recovery after photobleaching

Figure S16 shows predictions for photobleaching experiments for systems with several initial MT surface densities. An initially bleached circular patch increases in size and develops protrusions after the migration time $\tau_{\mathrm{N,max}}$. At high MT surface densities, the resulting bleaching patterns appear to be more compact and to have narrower protrusions than those for smaller MT surface fractions.

#### G. Single-microtubule dynamics: decomposing parallel-velocity distributions

It is difficult to pinpoint the origin of the asymmetry for parallel-velocity distributions that are obtained for all MTs in the system. Using the local polar order parameter, we are able to analyse the dynamics of MTs based on their local environments in more detail. Figure S17 decomposes the $v_{\|}$ distributions for $\phi = 0.3$ and $\tau^*$, shown in Fig. S14, into three distributions $v_{\|,A}$, $v_{\|,P}$, and $v_{\|,M}$ for MTs in antialigned, polar-aligned, and mixed environments, respectively, compare Fig. S3. For both, $p_a = 0.2$ and $p_a = 1.0$, the highest values of $v_{\|}$ are found for MTs in antialigned environments. The probability distributions in Figs. S17(a) and (b) show that the $v_{\|}$ distributions are symmetric for $p_a = 0.2$ and asymmetric and $p_a = 1.0$, for MTs in all three environments. However, the asymmetry of the distributions of MTs in antialigned environments is significantly more pronounced compared with MTs in polar-aligned and mixed environments. Furthermore, for both low and high MT surface fraction the active velocity overall shifts the $v_{\|}$ peak for antialigned MTs, but not for MTs in mixed and aligned environments.

In order to systematically compare the $v_{\|,M,P,A}$ distributions for $\tau^*$ we calculate the moments

$$m_r = \frac{1}{n}\sum_{i=1}^{n}(x_i - \langle x\rangle)^r \tag{3}$$

from the raw data so that we obtain

$$\text{mean} \quad \langle x\rangle = \frac{1}{n}\sum_{i=1}^{n} x_i\,, \qquad \text{variance} \quad m_2 = \frac{1}{n}\sum_{i=1}^{n}(x_i-\langle x\rangle)^2\,, \qquad \text{and skew} \quad \alpha_3 = \frac{m_3}{m_2^{3/2}}\,.$$

A generalised skewed distribution is defined as the product of the normal distribution and the cumulative distribution function,

$$f_s(x|\xi,\omega,\alpha) = \frac{2}{\omega}\Phi\left(\frac{x-\xi}{\omega}\right)\Psi\left(\alpha\left(\frac{x-\xi}{\omega}\right)\right). \tag{4}$$

with the cumulative distribution function

$$\Psi(x) = \int_{-\infty}^{x} dt\ \Phi(t) = \frac{1}{2}\left(1 + \text{erf}\left(\frac{x}{\sqrt{2}}\right)\right), \tag{5}$$

where erf is the error function and

$$\Phi(x) = \frac{1}{\sqrt{2\pi\omega^2}}e^{-(\frac{x-\xi}{\sqrt{2\omega^2}})^2}. \tag{6}$$

The moments of $f_s(x|\xi,\omega,\alpha)$ are the mean

$$\mu = \xi + \omega\delta\sqrt{\frac{2}{\pi}}, \tag{7}$$

the variance

$$\sigma^2 = \omega^2\left(1 - \frac{2\delta^2}{\pi}\right), \tag{8}$$

and the skew

$$\alpha_3 = \frac{(4-\pi)\left(\delta\sqrt{2/\pi}\right)^2}{2\left(1 - 2\delta^2/\pi\right)^{3/2}}, \tag{9}$$

where

$$\delta = \frac{\alpha}{\sqrt{1+\alpha^2}}. \tag{10}$$

Note that for $\alpha = 0$ we retrieve the original normal distribution with all its moments.

Figure S18(a), (b) and (c) show the values of $\mu$, $\omega$ and $\alpha$ as calculated by the procedures outlined above, best fits were obtained with $\alpha = 0$ for $\mu(v_{\|,M,P})$ and $\alpha > 0$ for $\mu(v_{\|,A})$. In particular, the mean $\mu(v_{\|,A})$ increases much stronger with $p_a$ than $\mu(v_{\|,P})$ and $\mu(v_{\|,M})$. Polar-aligned MTs are slowest, and a non-zero $\mu(v_{\|,P})$ is likely due to dragging of polar-aligned MTs due to streaming of antialigned MTs by friction between MTs and collisions with antialigned MTs [8]. Figure S18(b) shows that the variance increases both with $\psi_i$ and $p_a$, and decreases with $\phi$. The variance is highest and increases strongest with $p_\mathrm{a}$ for MTs in antialigned environments. Figure S18(c) shows that the skews increase with activity. The distributions of MTs in antialigned environments have smaller skews than of MTs in polar-aligned and mixed environments. The asymmetric distributions for MTs in mixed and polar-aligned environments reflects both passive motion (which is symmetric) and being dragged along with propelled MTs by friction. Figure S18(d) shows the differences in distributions of $v_\parallel$ due to increasing motor activity for $\phi = 0.3$. We quantify these differences, by computing moments of the $v_\parallel$ distributions for varying $\phi$ and $p_\mathrm{a}$ for the three categories of MT environments.

Figure S19 provides data for parallel MT-velocity distribtions for all MTs for various MR surface fractions, analogously to Fig. 6 in the main text.

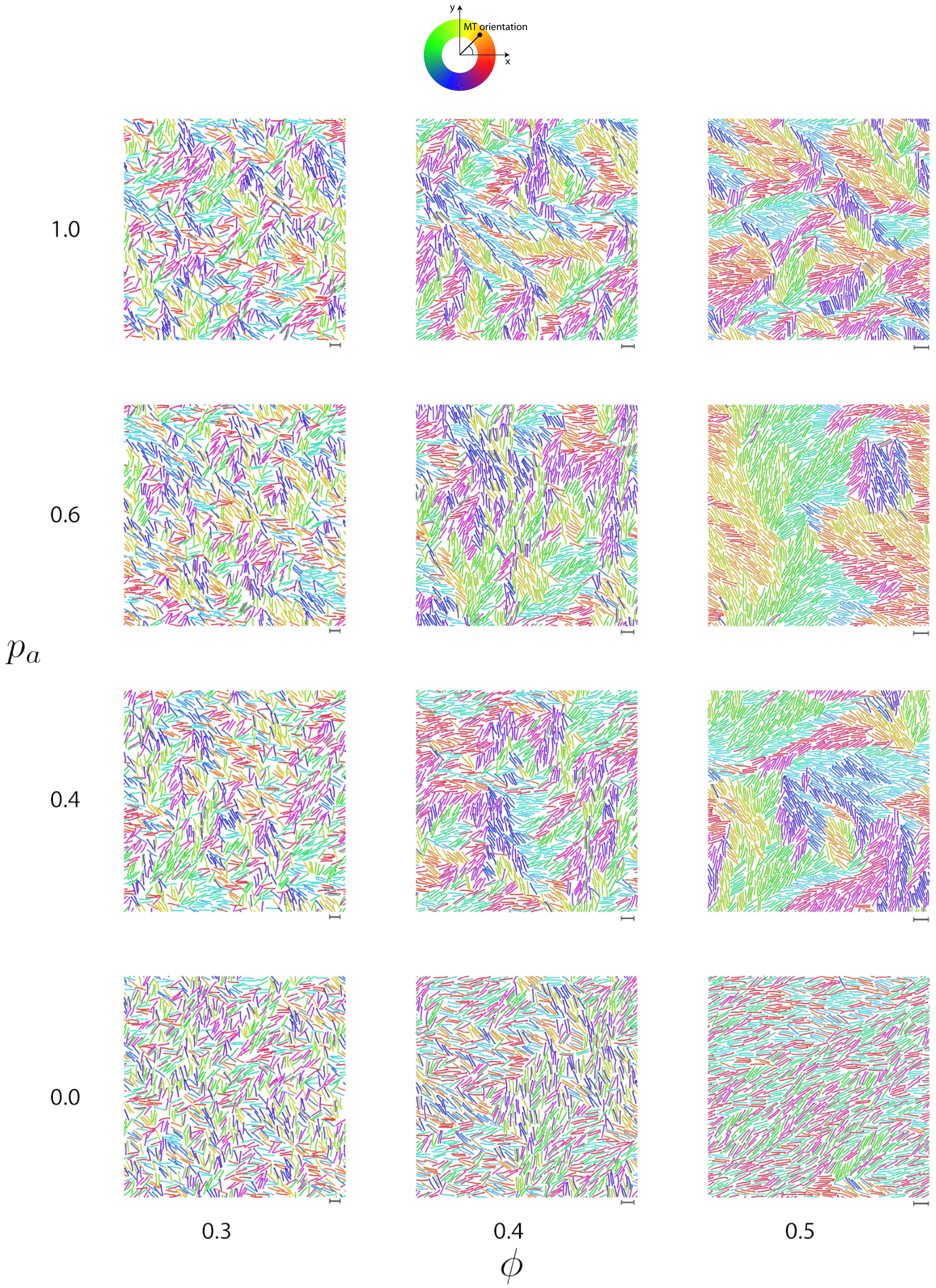


FIG. S2. Simulation snapshots at steady-state for systems with varying antialigned motor probabilities $0.0 < p_{\rm a} \leq 1.0$, and MT surface fractions $0.3 < \phi < 0.5$. The surface fractions of MTs are varied by changing the size of the periodic box, while keeping the number of MTs constant. The scale bars that correspond to the length of a MT is indicated in the bottom right of each simulation frame. The colours represent the orientation of the polar MTs with respect to the system reference frame according to the colour wheel above.

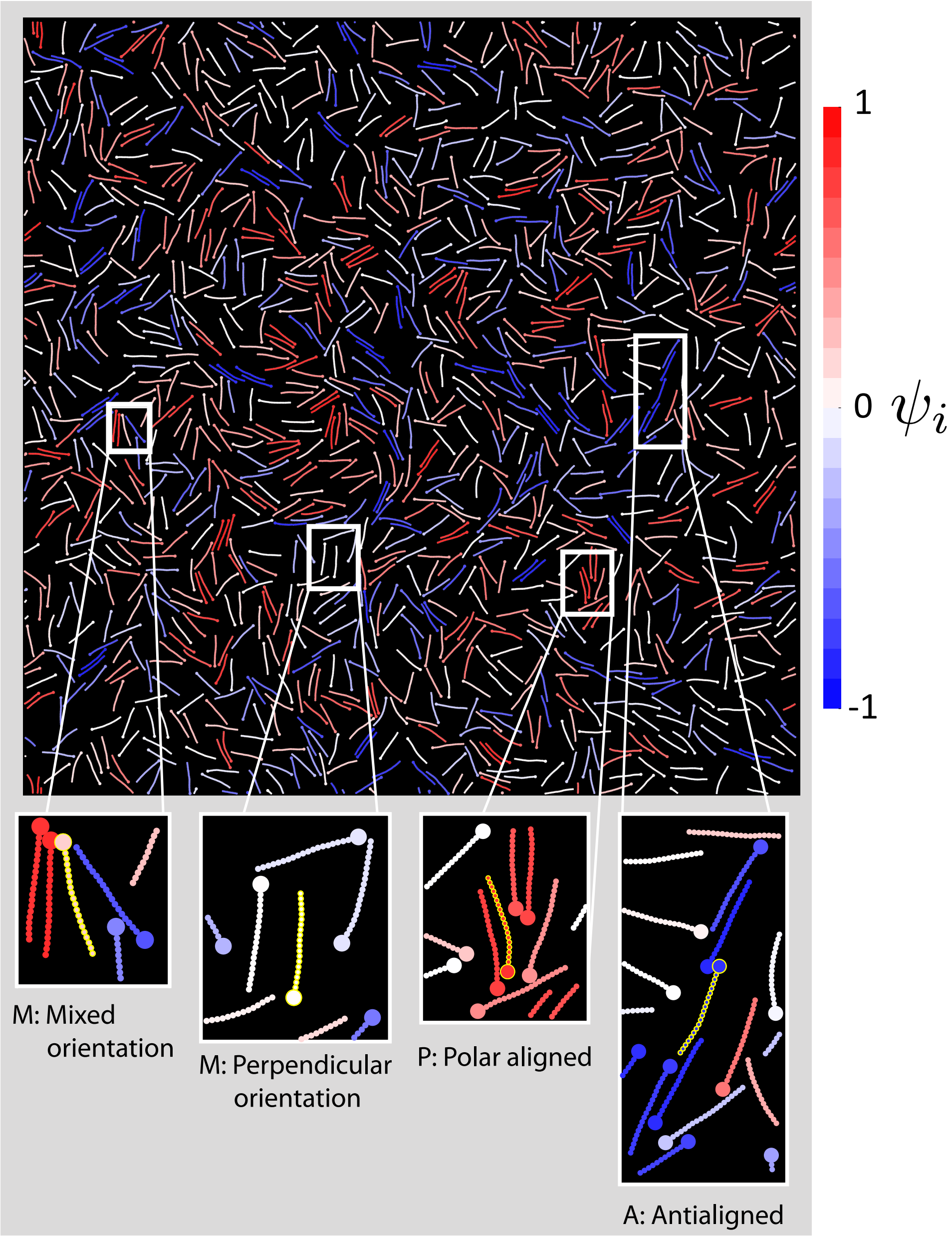


FIG. S3. MTs coloured based on their local polar order parameter $\psi_i$ for $p_a = 1.0$, $\phi = 1.0$. The colour corresponding to $-1 < \psi_i < 1$ is given on the right. Zoomed in illustrations of MTs show examples of MTs in the three $\psi_i$ categories distinguished in Fig. S4. The MT in question is highlighted in yellow in the zoomed in graphics. (M) $\psi_i \approx 0$ values can occur either when MTs are perpendicularly oriented with respect to its surrounding or when MTs have neighbours which are both polar-aligned and antialigned. (P) $\psi_i > 0.5$ occurs when MTs have neighbours which are mostly polar-aligned. (A) $\psi_i < -0.5$ occurs when MTs have neighbours which are mostly antialigned.

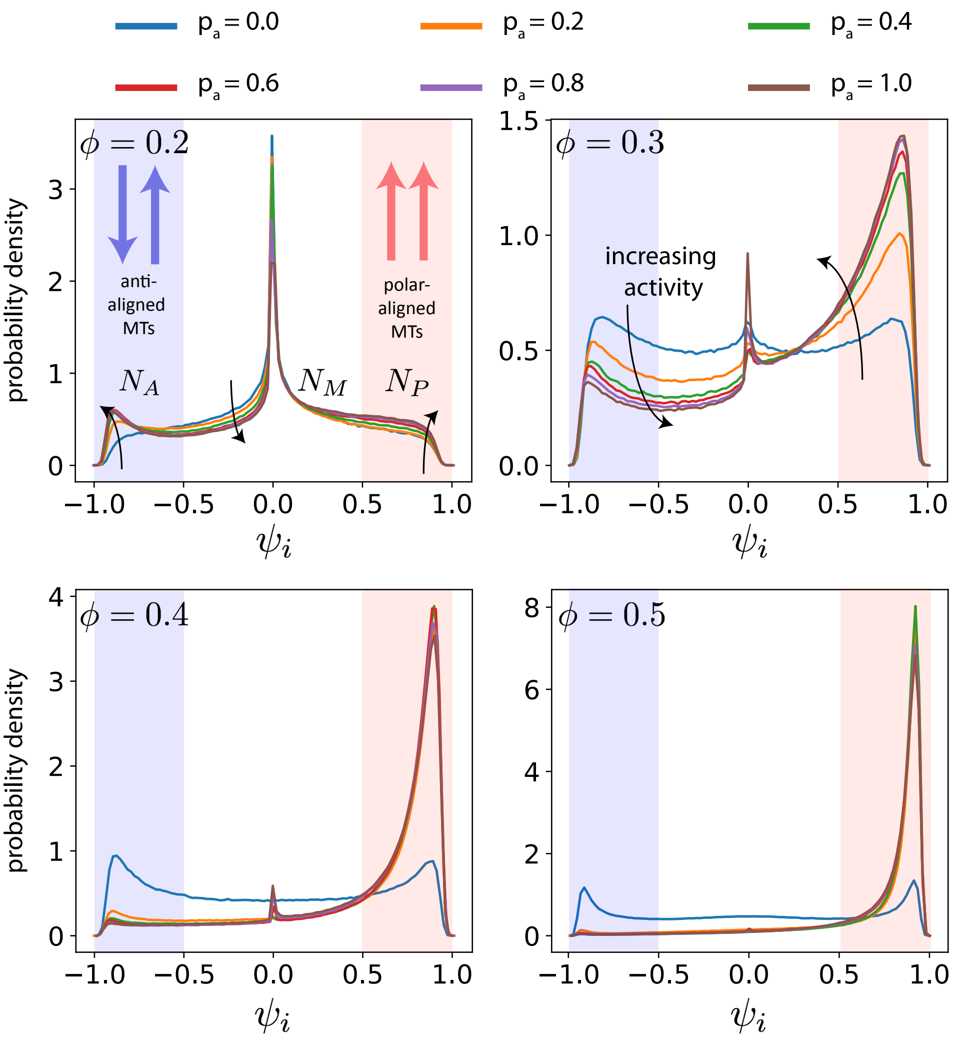


FIG. S4. Probability densities of MTs' local polar order parameter $\psi_i$ as function of $p_a$ and $\phi$. The arrows are guides, which indicate the directions in which activity increases. In the following sections MTs are distinguished based on their relative orientation to their local environment. $A$, $M$ and $P$ indicate MTs that have local polar order corresponding to $\psi_i \leq -0.5$ (green), $-0.5 < \psi_i < 0.5$ (white), $\psi_i \geq 0.5$ (blue), respectively. $n_A$, $n_m$ and $n_P$ are the number of MTs which are in antialigned, perpendicular and polar-aligned environments respectively. Note that the scale of the ordinate axis is different for each surface fraction.

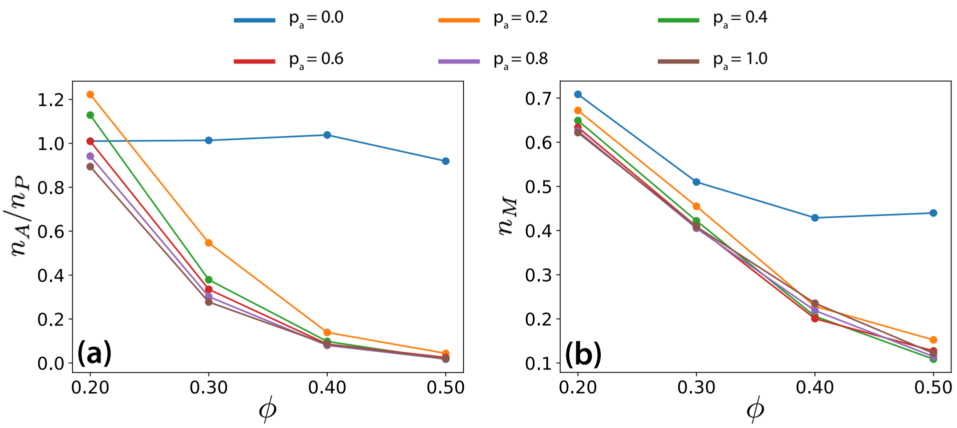


FIG. S5. Ratios of MT populations in (a) antialigned to polar-aligned environments, and (b) perpendicular environments to total number of MTs, for various MT surface fractions $\phi$ and antialigned motor probabilities $p_a$.

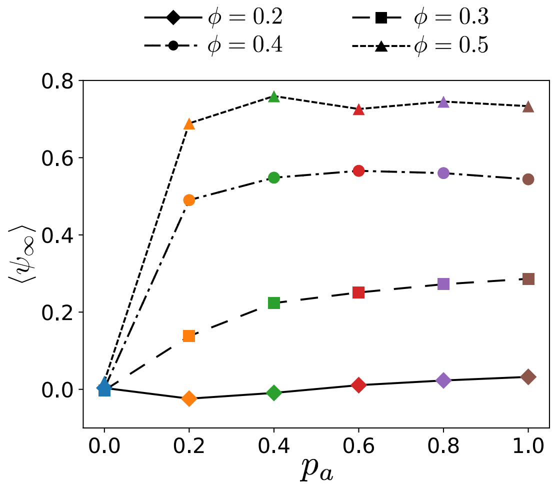


FIG. S6. Mean local polar order parameter of MTs at long times, $\langle \psi_\infty \rangle$, for various $\phi$ and $p_\mathrm{a}$.

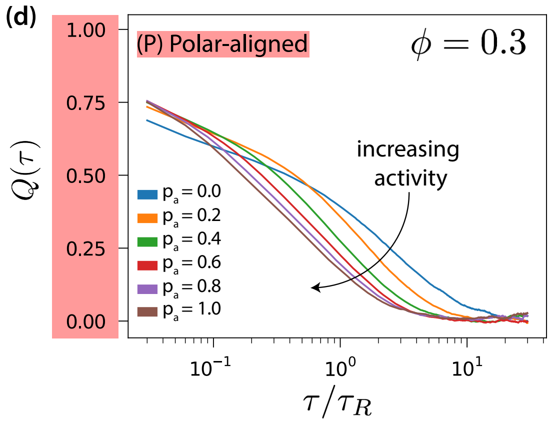


FIG. S7. Deviation from local polar order $Q(\tau)$ for various $p_a$ for aligned MTs.

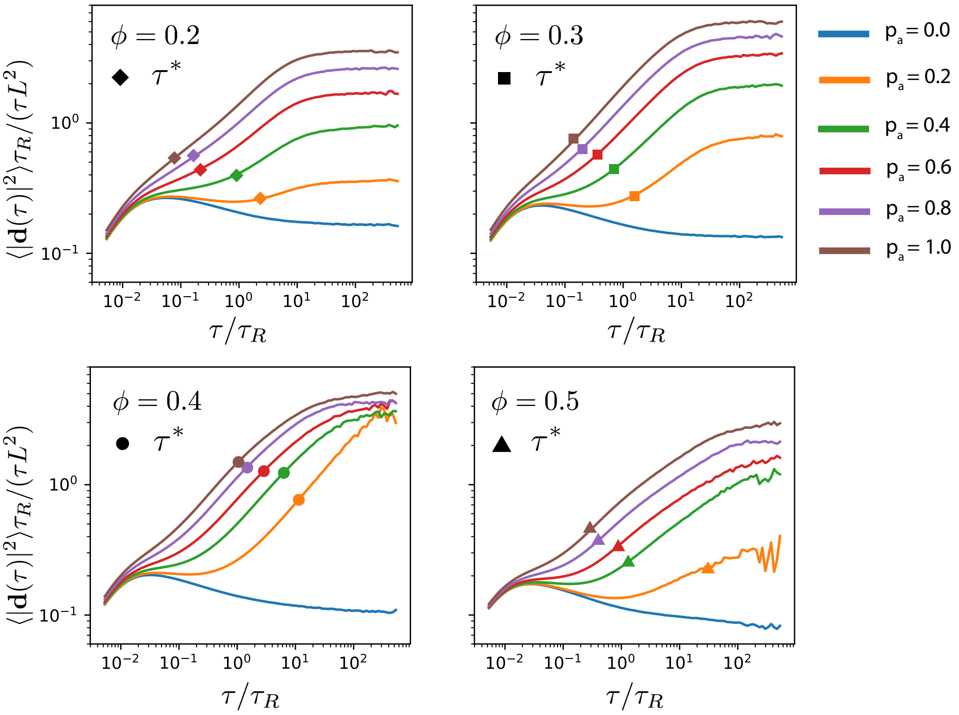


FIG. S8. Translational MSD/Lag time for various activities, $p_{\rm a}$, and surface fractions, $\phi$. MSD is normalised using $L^2/\tau_R$, and lag time, $\tau$, is normalised using $\tau_R$. The symbols on the plots indicate the time-scale at which parallel velocity is maximally skewed due to active forces.

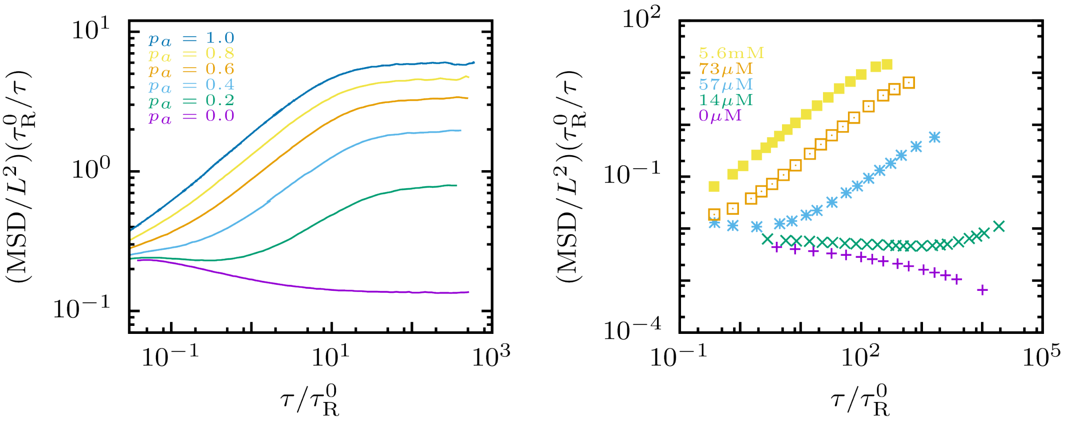


FIG. S9. **a** Normalized MSD curves from our simulations for several motor probabilities $p_a$. **b** Normalized MSD curves as a function of lag time for experiments from Ref.[7] for selected ATP concentrations. Here $L$ is the filament length, $\tau^0_R$ the single filament rotation time and $\tau$ the lag time.

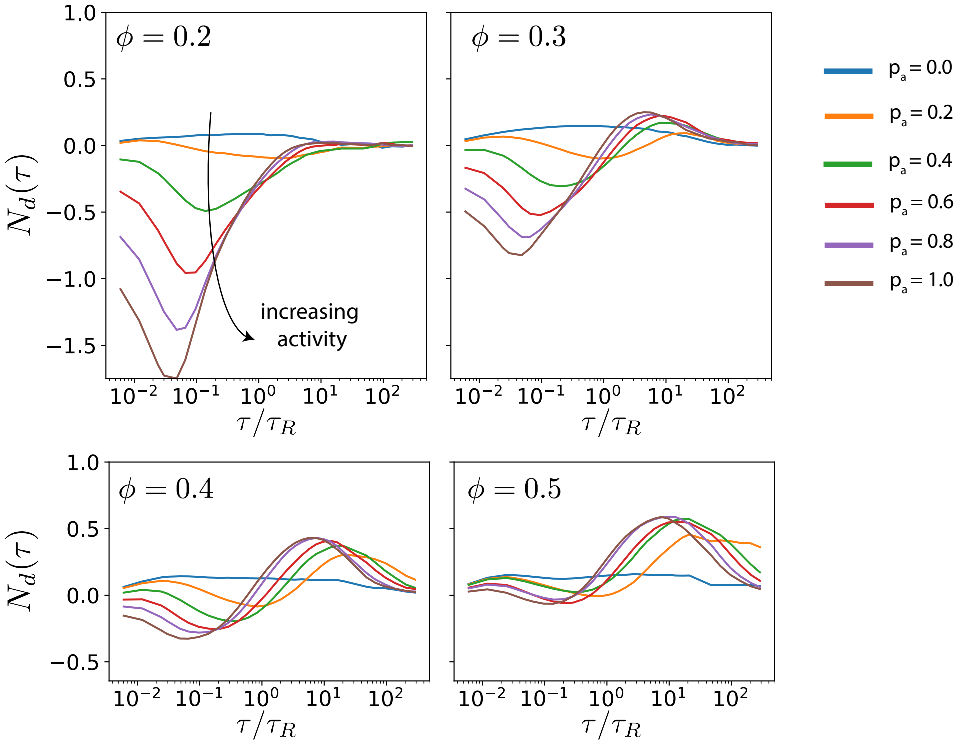


FIG. S10. Neighbour displacement correlation function $N_{\mathrm{d}}(\tau)$ for different $\phi$ and $p_{\mathrm{a}}$. The time at which the minimum and maximum of $N_{\mathrm{d}}(\tau)$ occurs are $\tau_{N,\mathrm{min}}$ and $\tau_{N,\mathrm{max}}$ respectively.

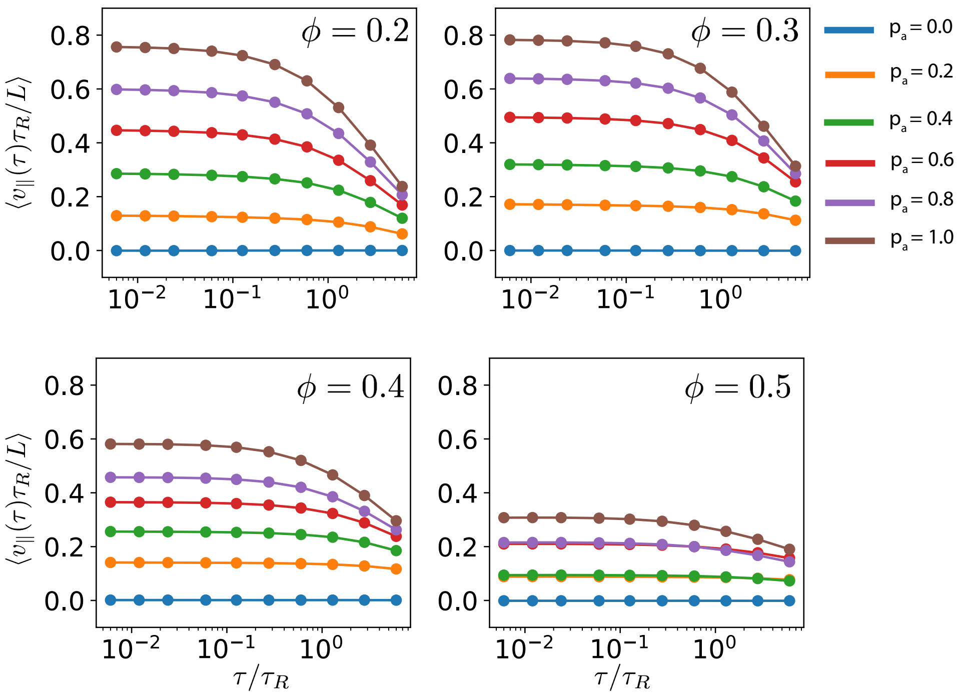


FIG. S11. Parallel velocity $v_\parallel$ as a function of $\tau$ for different area fractions and different activities.

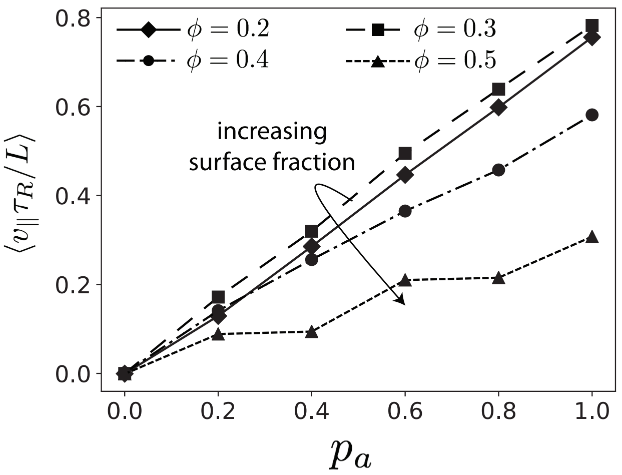


FIG. S12. Parallel velocity $v_{\|}$ extrapolated to $\tau = 0$ as a function of $p_{\rm a}$ for different area fractions.

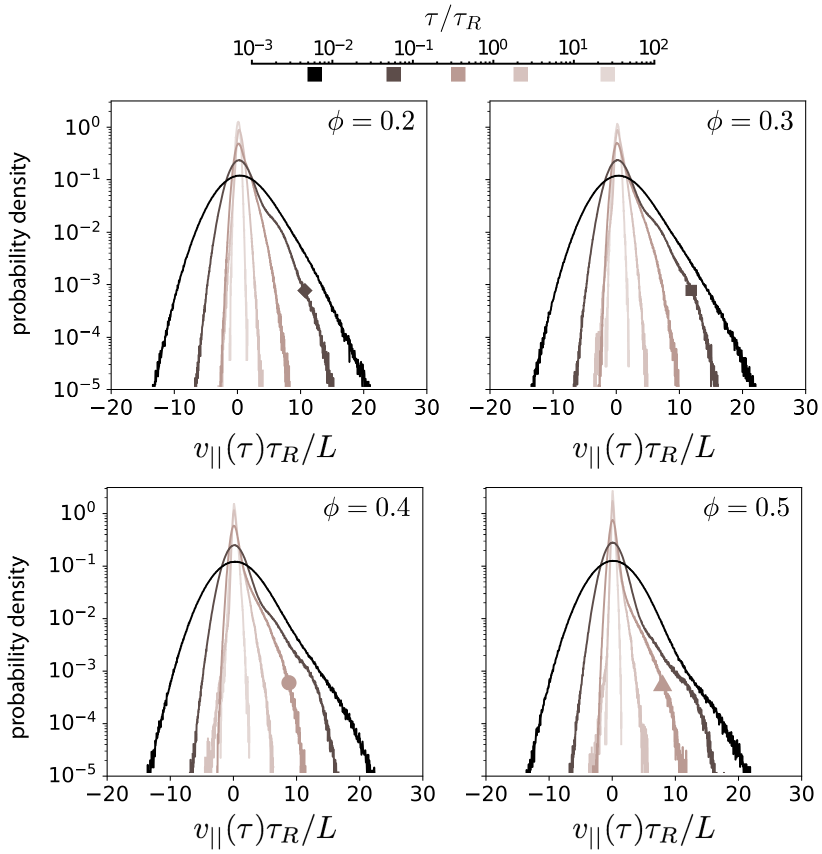


FIG. S13. Histogram of $v_{\|}$ for various surface fractions $\phi$ and five time windows $\tau$. The darkness of the curve represents the time window used to measure the parallel velocity. The darkest-coloured curve represents the parallel velocity obtained for the shortest time window, and the lightest-coloured curve is obtained from the longest time window. The box symbols on the curves correspond to the displacement distribution that is closest to the distribution which is most skewed.

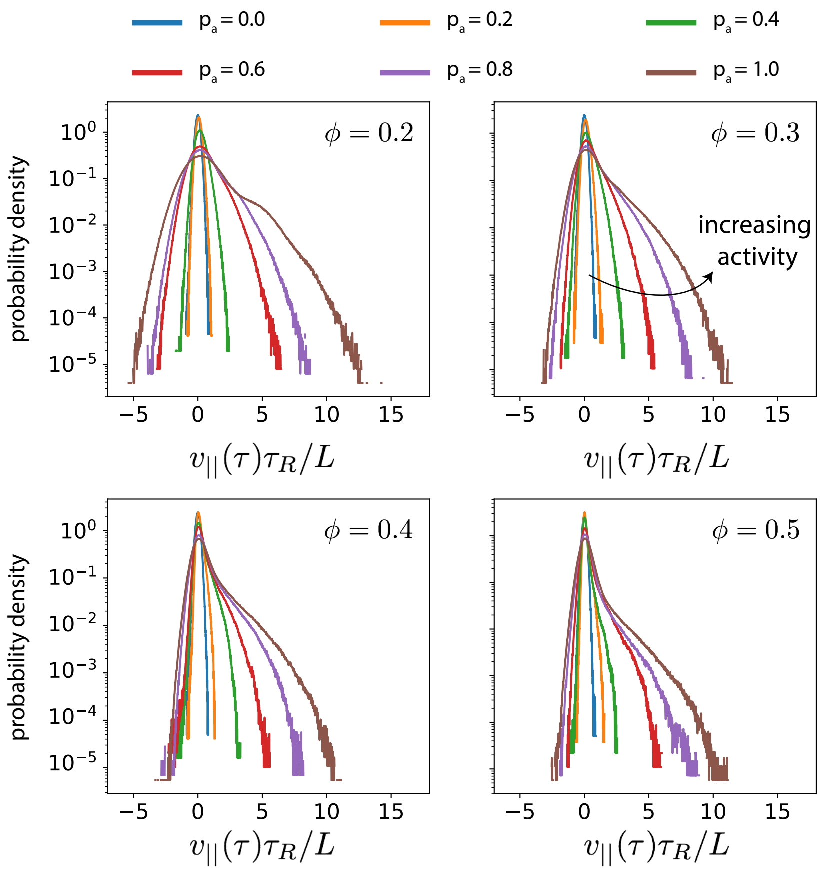


FIG. S14. Histogram of $v_{\|}(\tau^*)$ for various $p_a$ and $\phi$. The duration of the time window corresponds to the maximal skew, see Fig. S13. This indicates the structure of the velocity distribution when the skew is maximal. The ordinate axis is log scaled to show the deviation of the distribution from a Gaussian, which would appear as a symmetric inverted parabola.

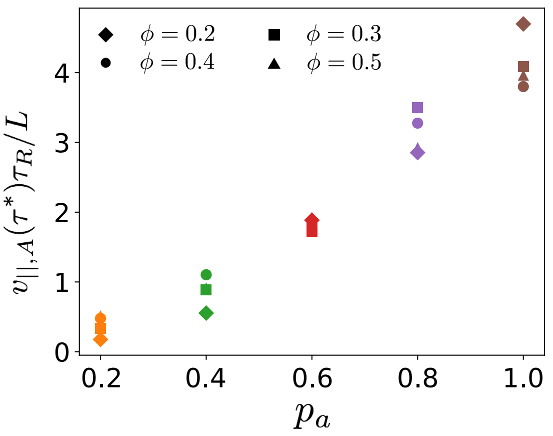


FIG. S15. Maximum parallel MT velocities $v_{\|,A}(\tau^*)$ as function of $p_a$ for various MT surface fractions $\phi$.

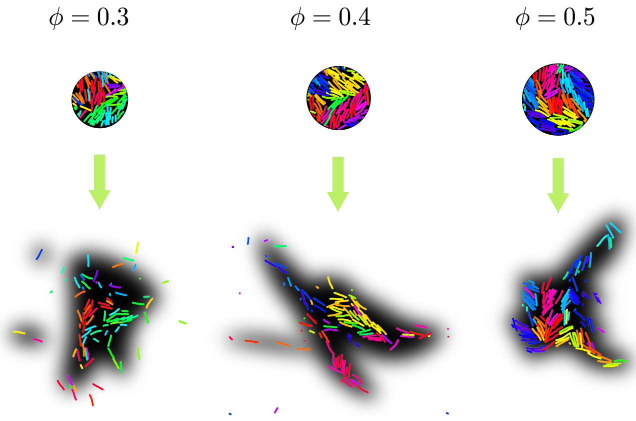


FIG. S16. Predictions for a photobleaching experiment with $\phi = 0.3, 0.4$ and $0.5$. MTs retain the orientation colour from when they were tagged at $t = 0$. The black shadow shows our predictions for photobleaching experiments at time $\tau_{\rm N,max}$ after bleaching a circular patch.

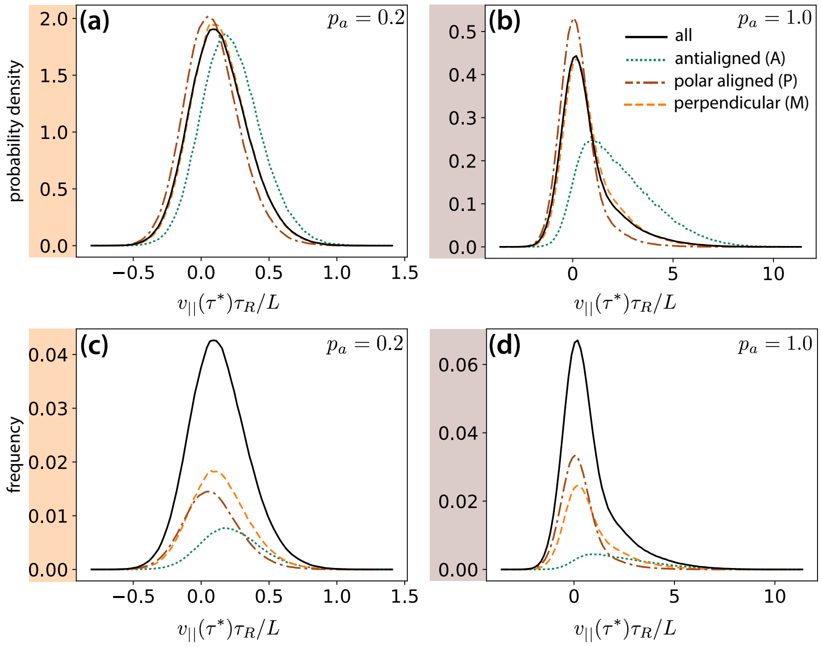


FIG. S17. MT parallel velocity distributions $v_\parallel$ for a time window of duration $\tau^*$ and $\phi = 0.3$, decomposed based on MT environments (A, M, P) determined by their local polar order parameter, $\psi_i$, see Fig. S4. **(a)** and **(b)** show probability density histograms of $v_\parallel(\tau^*)$ for $p_{\rm a} = 0.2$ and 1.0, respectively. **(c)** and **(d)** show frequencies of occurrence of $v_\parallel(\tau^*)$ for $p_{\rm a} = 0.2$ and 1.0, respectively. The sum of the decomposed curves in (c) and (d) gives the solid curve shown.

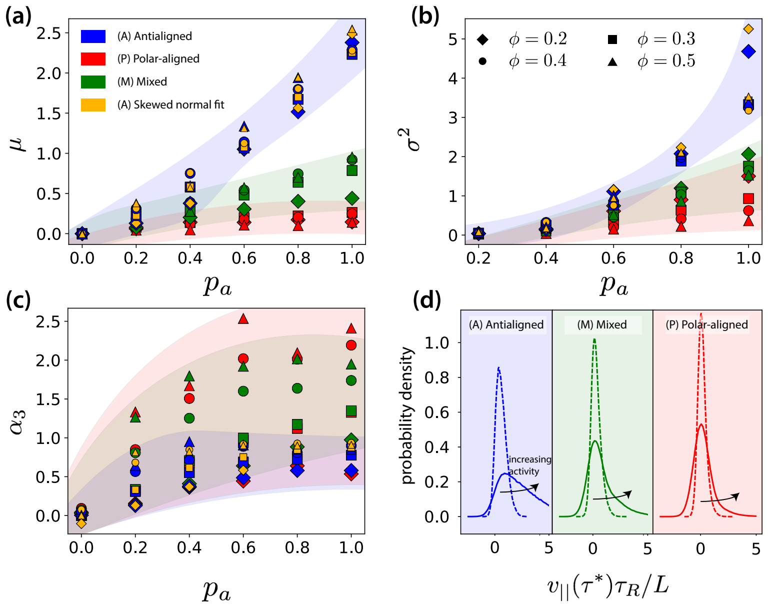


FIG. S18. First three moments, **(a)** mean, **(b)** variance, **(c)** skew of the $v_{\|}$ distribution for a time window of duration $\tau^*$ for MTs in (A) antialigned, ($\psi_i < -0.5$, blue), (P) polar-aligned ($\psi_i > 0.5$, red), and (M) mixed ($|\psi_i| \leq 0.5$, green) environments, for different $\phi$ and $p_a$. The blue, red and green markers indicate moment calculated from raw data. The yellow markers are obtained from calculating moments from fits to the antialigned parallel MT velocity distribution $v_{\|,A}$. **(d)** Example of differences in structures of $v_{\|}$ distributions due to increasing activity from $p_a = 0.4$ (dotted line) to $p_a = 1.0$ (solid line) for $\phi = 0.3$, for A, P and M categories of MT environment. Compare also with Fig. S16.)

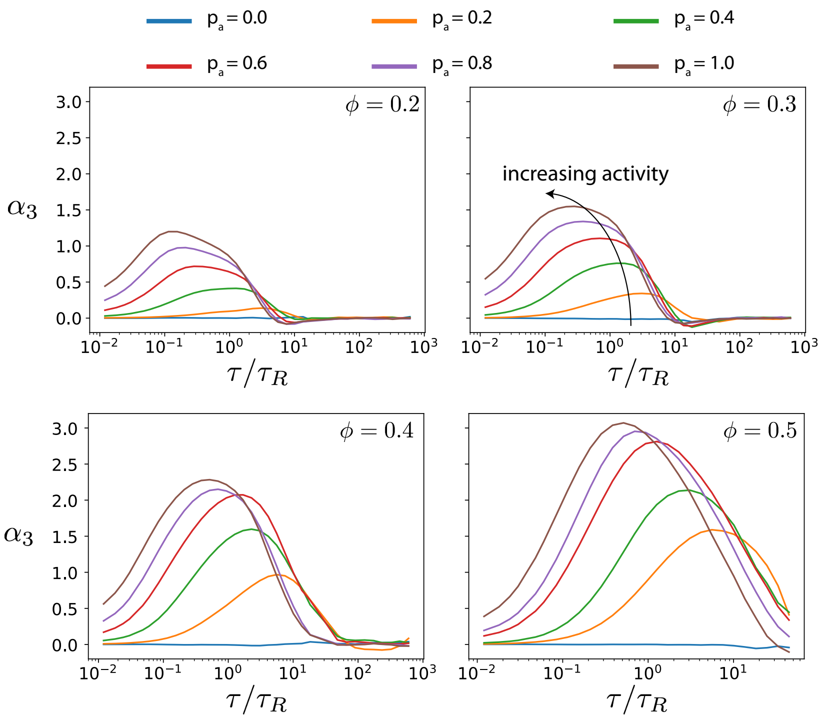


FIG. S19. Skews $\alpha_3$ of $v_{\|}$ distributions (Fig. S13) computed as a function of lag times for different $p_a$ and $\phi$. The probability distributions that correspond to the maximal skew are shown in Fig. S13 together with distributions for few other lag times. The ordinate scale is the same for comparison of overall skew between different MT surface fractions.

[1] A. Ravichandran, G. A. Vliegenthart, G. Saggiorato, T. Auth, and G. Gompper, Biophys. J. **113**, 1121 (2017).
[2] M. A. Bates and D. Frenkel, J. Chem. Phys. **112**, 10034 (2000).
[3] This includes also MTs in low surface-fraction systems that do not have contacts with neighbouring MTs or that are perpendicularly oriented. However, such instances are rare even at the lowest surface fraction that we studied, $\phi = 0.2$.
[4] J. R. Howse, R. A. L. Jones, A. J. Ryan, T. Gough, R. Vafabakhsh, and R. Golestanian, Phys. Rev. Lett. **99**, 048102 (2007).
[5] J. Elgeti, M. Cates, and D. Marenduzzo, Soft Matter **7**, 3177 (2011).
[6] R. E. Isele-Holder, J. Elgeti, and G. Gompper, Soft Matter **11**, 7181 (2015).
[7] T. Sanchez, D. T. Chen, S. J. DeCamp, M. Heymann, and Z. Dogic, Nature **491**, 431 (2012).
[8] A non-zero $\mu(v_{\|,P})$ can also occur from a change of local polar order of an MT during measurement time.
